# Leveraging Foundation Models for the Characterisation of Small RNA Properties

**DOI:** 10.64898/2026.02.08.704350

**Authors:** Shivprasad Jamdade, Coyun Oh, Heba Sailem

## Abstract

Small interfering RNAs (siRNAs) provide a promising therapeutic approach capable of selectively silencing disease-associated genes; however, achieving high efficacy and specificity while minimising off-target effects remains a significant challenge. Endogenous small RNAs, such as microRNAs (miRNAs) and PIWI-interacting RNAs (piRNAs), exhibit structural features supporting their functions and are biocompatible. Recent advances in RNA foundation models, such as RNA-FM, enable large-scale learning of sequence and structural representations of RNA sequences, offering a powerful framework for studying small RNA functions. Here, we leverage RNA-FM model alongside interpretable biological features to systematically compare endogenous small RNAs (miRNAs and piRNAs) with synthetic siRNAs. Biological features highlighted class-specific patterns: piRNAs showed significantly higher GC content and melting temperature than miRNAs and siRNAs, suggesting higher stability. Importantly, we mapped RNA-FM embeddings to interpretable features to better understand deep learning outputs and facilitate effective extraction of functionally relevant information. To support predictive and comparative analyses of small RNAs, we implemented these functionalities in RNAExplorer (www.rnaexplorer.com), a web-based application that allows analysing and visualising small RNA features interactively. Together, our integrative analysis provides a framework for understanding small RNA biology and improving siRNA therapeutic design strategies.

**Graphical Abstract:** 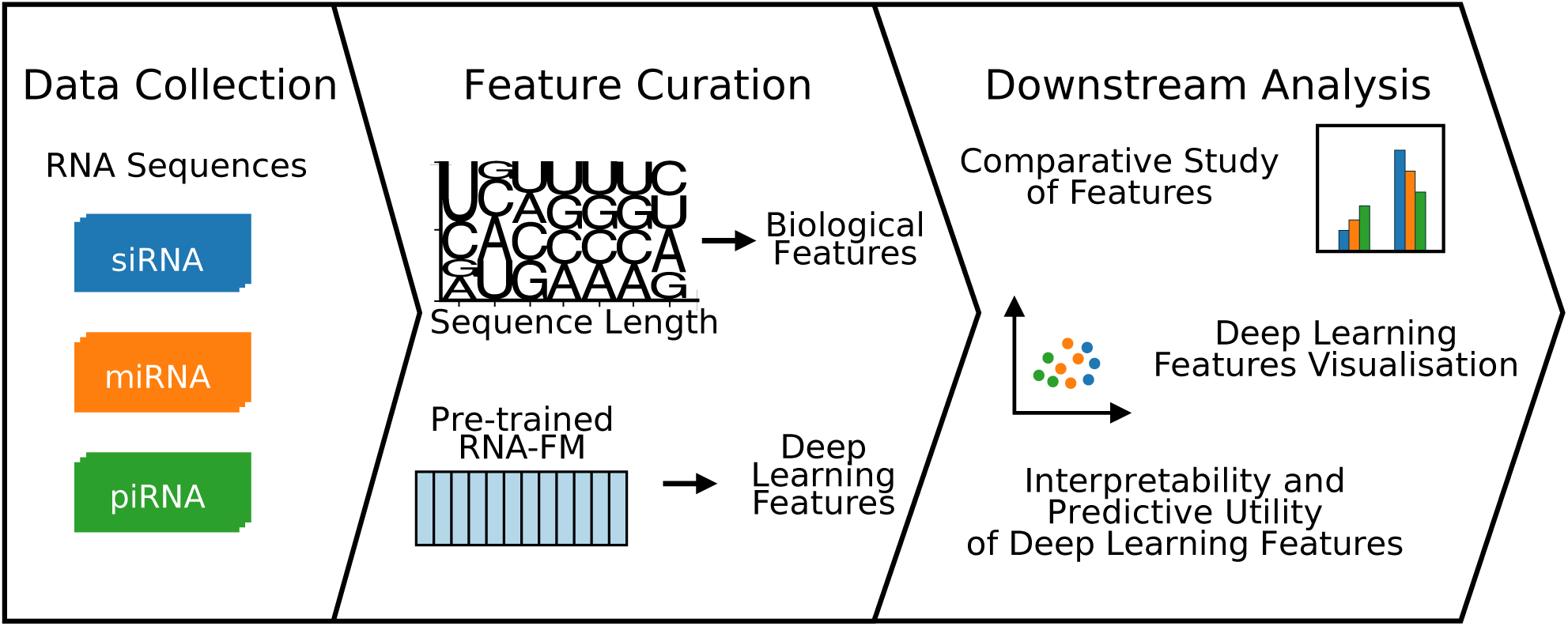

## INTRODUCTION

Small interfering RNAs (siRNAs) have emerged as powerful tools for selectively silencing disease-associated genes by exploiting the conserved RNA interference machinery present in cells. While inspired by natural RNA regulatory pathways, the development of therapeutic siRNAs involves multiple critical stages, each presenting distinct challenges (1). First, sequence design must ensure effective target recognition while minimising off-target effects, as even partial complementarity can lead to off-target gene silencing (2). In particular, the siRNA seed region plays a key role in triggering off-target effects, as short matches to unintended transcripts can cause microRNA-like gene silencing (2,3). Another challenge is that delivery to target tissues and cells remains a major hurdle (4). Finally, preclinical and clinical testing frequently reveal unanticipated toxicity, immune responses, or insufficient efficacy, leading to the failure of many candidate siRNAs in clinical trials (5). These challenges motivate the development of approaches that systematically identify features that contribute to siRNA efficacy and cellular compatibility (6). Indeed, many studies focused on studying features contributing to siRNA efficacy, albeit mostly based on in vitro assays (7). However, there is limited large-scale comparison with the structure of endogenous RNA that might provide valuable insights into biocompatibility (8).

Endogenous small non-coding RNAs are essential regulators of gene expression and genome stability in eukaryotes. In animals, microRNAs (miRNAs), typically 22 nucleotides in length, control post-transcriptional repression of target mRNAs via Argonaute proteins, modulating diverse developmental and physiological processes (3). Small interfering RNAs (siRNAs), often 20–23 nucleotides long, predominantly arise from double-stranded RNA precursors and trigger sequence-specific mRNA degradation through the RNA interference pathway (9). PIWI-interacting RNAs (piRNAs, 24–31 nucleotides long) are a distinct class expressed primarily in germline cells. piRNAs associate with PIWI proteins, a subfamily of Argonaute proteins named P-element-induced wimpy testis, to silence transposable elements and protect genome integrity (10). These RNA classes differ in their targets and biogenesis pathways (Drosha/Dicer processing for miRNAs and siRNAs versus Dicer-independent piRNA biogenesis in germline), yet all function via the Argonaute/PIWI-family effector complexes (11).

Studying the sequence and structural features that distinguish siRNAs, miRNAs, and piRNAs can provide valuable insights into their differences and similarities and how this can support their efficacies and functions. Prior studies have identified characteristic motifs or biases; for example, many miRNAs and piRNAs show a preference for 5’-uridine, which influences Argonaute/PIWI loading (10,11). However, a large-scale comparative analysis of small RNA features that can provide deeper insight into their structural features has yet to be explored.

Traditional analyses often focus on individual features such as nucleotide composition or secondary structure stability that are known to affect siRNA efficacy. Recent advances in deep learning enable more holistic representations of RNA sequences. For instance, RNA-FM is a large-scale RNA foundation model trained on more than 23 million RNA sequences in a self-supervised manner. It utilises a transformer encoder pretrained through masked language modelling, in which selected nucleotides are hidden and predicted from the surrounding sequence context. Through this process, RNA-FM learns high-dimensional numerical representations of RNA sequences, known as embeddings, without requiring sequence annotations (12). Such embeddings have demonstrated utility in predicting sequence efficacy as well as RNA secondary and tertiary structure (12). We hypothesised that RNA-FM embeddings would support predictive tasks and provide insights into the distinguishing characteristics of siRNAs, miRNAs, and piRNAs, complementing the analysis of interpretable sequence features.

Although individual biochemical properties of small RNAs have been studied extensively, a systematic comparison of multiple compositional, positional, structural and thermodynamic properties across siRNAs, miRNAs and piRNAs, together with an evaluation of the utility of RNA foundation-model representations for interrogating these properties, has been lacking. Here, we address this gap by establishing a quantitative reference landscape across these classes and translating the resulting framework into RNAExplorer, an accessible resource for comparative small-RNA analysis.

In this work, we analyse more than 48,000 small RNA sequences, integrating interpretable features and RNA-FM feature embeddings to determine the feature space of various small RNAs. We compiled a large dataset of siRNAs, miRNAs, and piRNAs with their defined features, including sequence composition, Shannon entropy of nucleotide distribution, AU and GC skew, and thermodynamic stability. Using these features, we compare the distributions and correlations across the three RNA classes. We further leverage the pre-trained RNA-FM model to obtain embeddings for each small RNA and visualise their relationships in a reduced-dimensional space. Our study highlights both expected and novel patterns. For instance, the extreme 5’-uridine bias of piRNAs, the uridine-rich preference of siRNAs, and the modest enrichment of uridine in miRNAs are all reflected in the RNA-FM embedding space. Most importantly, we developed a user-friendly web application, RNAExplorer, to allow a wide range of scientists to explore sequences, integrate them within the broader RNA landscape, and analyse both interpretable and deep learning features. By combining traditional feature analysis with deep learning representations, our study provides a comprehensive view of the small RNA sequence landscape and opens new avenues for classifying and understanding small RNAs.

## MATERIALS AND METHODS

### Experimental and Technical Design

Figure 1 illustrates the workflow of our methodology, from dataset preparation and feature curation to downstream tasks.

**Figure 1.**
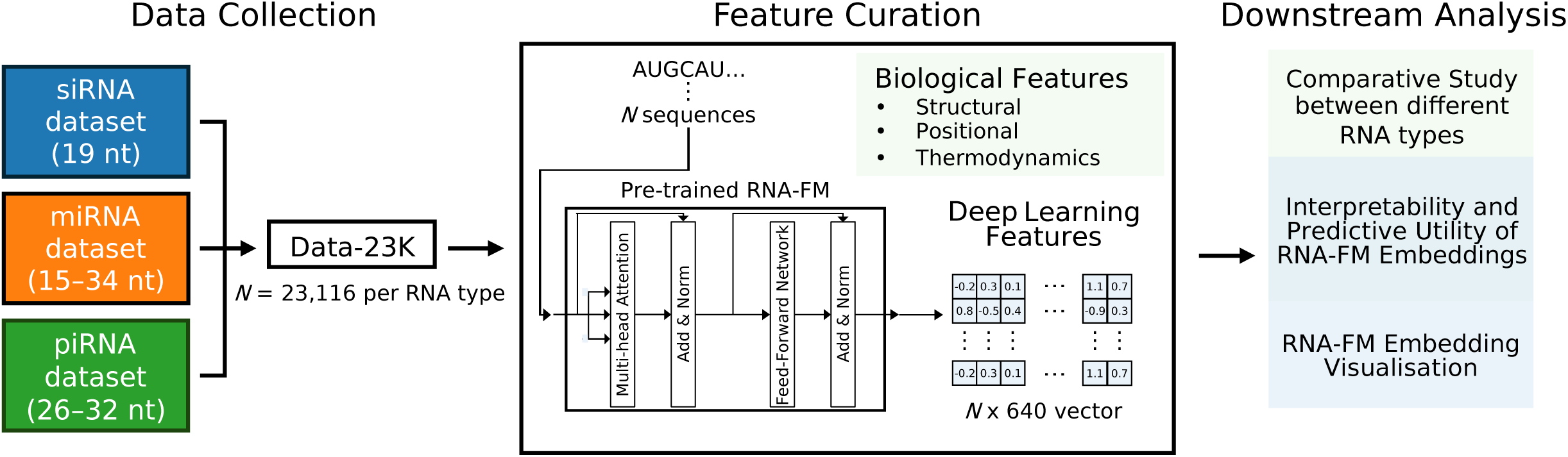
Workflow of the methodology in this paper.

### Datasets

Antisense and sense siRNA sequences were sourced from several previous studies (13–15) (Table 1). To ensure consistent antisense strand orientation, siRNA sequences with sense strand orientation from Takayuki (14) and siRNA screen (15) datasets were reverse complemented. When present, 3’ overhangs were removed, so that the sequences are directly comparable with miRNA and piRNA sequences. We obtained miRNA sequences from miRBase, the primary catalogue for published miRNA sequences (16,17). The piRNA dataset was obtained from the human small ncRNA database, DASHR2-GEO (18). Any precursors were excluded, and only unique mature RNA sequences were used to avoid data redundancy caused by isoforms or different species.

**Table 1.** Details of each dataset used in this paper, with nt indicating nucleotide.

| Dataset | Number of Mature Sequences | Number of Unique Sequences | Sequence Length (nt) | Species | Source |
| --- | --- | --- | --- | --- | --- |
| siRNA | 2,431 | 2,431 | 19 | Homo Sapiens | Huesken (13) |
|  | 702 | 702 | 19 |  | Takayuki (14) |
|  | 71,625 | 71,557 | 19 |  | siRNA screen (15) |
| miRNA | 48,885<br>(2,656 Homo Sapiens) | 30,721<br>(2,632 Homo Sapiens) | 15–34<br>(16–28 for Homo Sapiens) | 271 species, including Homo Sapiens | (16,17) |
| piRNA | 50,397 | 23,116 | 26–32 | Homo Sapiens | (18) |
| Total | 174,040 | 128,515* | 15–34 |  |  |

\* Total number of sequences after removing duplicate siRNA sequences from the original total of 128,527 sequences. One sequence was annotated as a mature miRNA from both *Mus musculus* (MIMAT0031410) and *Cricetulus griseus* (MIMAT0041163), and as a *Homo Sapiens* piRNA (piR-54381), and is therefore retained in our downstream analysis.

To ensure balanced comparisons and prevent dataset size bias, three subsets were created from all five datasets. Data-Whole includes total unique sequences from the three subsets (128,515). Data-23K was generated by randomly selecting 23,116 sequences from each RNA type to provide a large, balanced dataset for downstream analyses. Because human miRNAs had only 2,632 unique sequences, additional miRNA sequences from other species were included after confirming that distributions of sequence features in the smaller, human-only subset (Data-3K, 2,632 sequences per RNA type) were consistent with those in the larger Data-23K. Specifically, Pearson correlation analysis between interpretable features and RNA-FM embedding dimensions across these three subsets showed that Data-23K closely represents both Data-Whole (*r ≈* 0.9989-0.9999) and Data-3K (*r ≈* 0.9322-0.9941), justifying the use of miRNA from other species for subsequent analyses to mitigate sample size bias (Supplementary Fig. S1).

### Selection and Computation of Interpretable Features

For each RNA sequence, a set of biologically interpretable features was extracted to quantify thermodynamic stability, sequence composition, and structural complexity (Table 2). Thermodynamic descriptors included Gibbs free energy (ΔG) and Minimum Free Energy (MFE), both computed using the RNAfold program from the ViennaRNA package (v2.4) under standard conditions (37°C, 1 M NaCl; M indicate molar concentration) (19). These parameters represent the predicted secondary structure stability of each RNA based on minimum-energy folding models. Additionally, the melting temperature (Tm) was calculated with Biopython’s Melting Temperature module via the nearest-neighbour method.

**Table 2.**
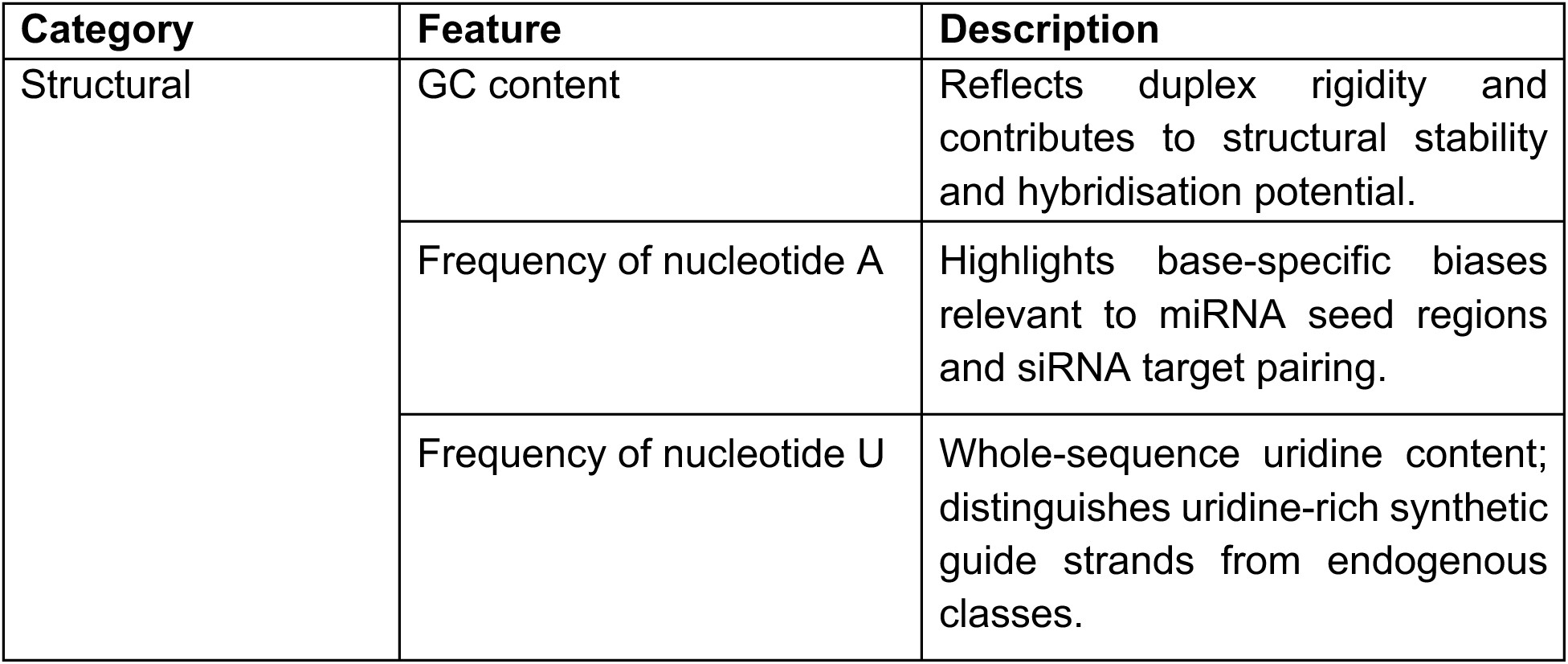
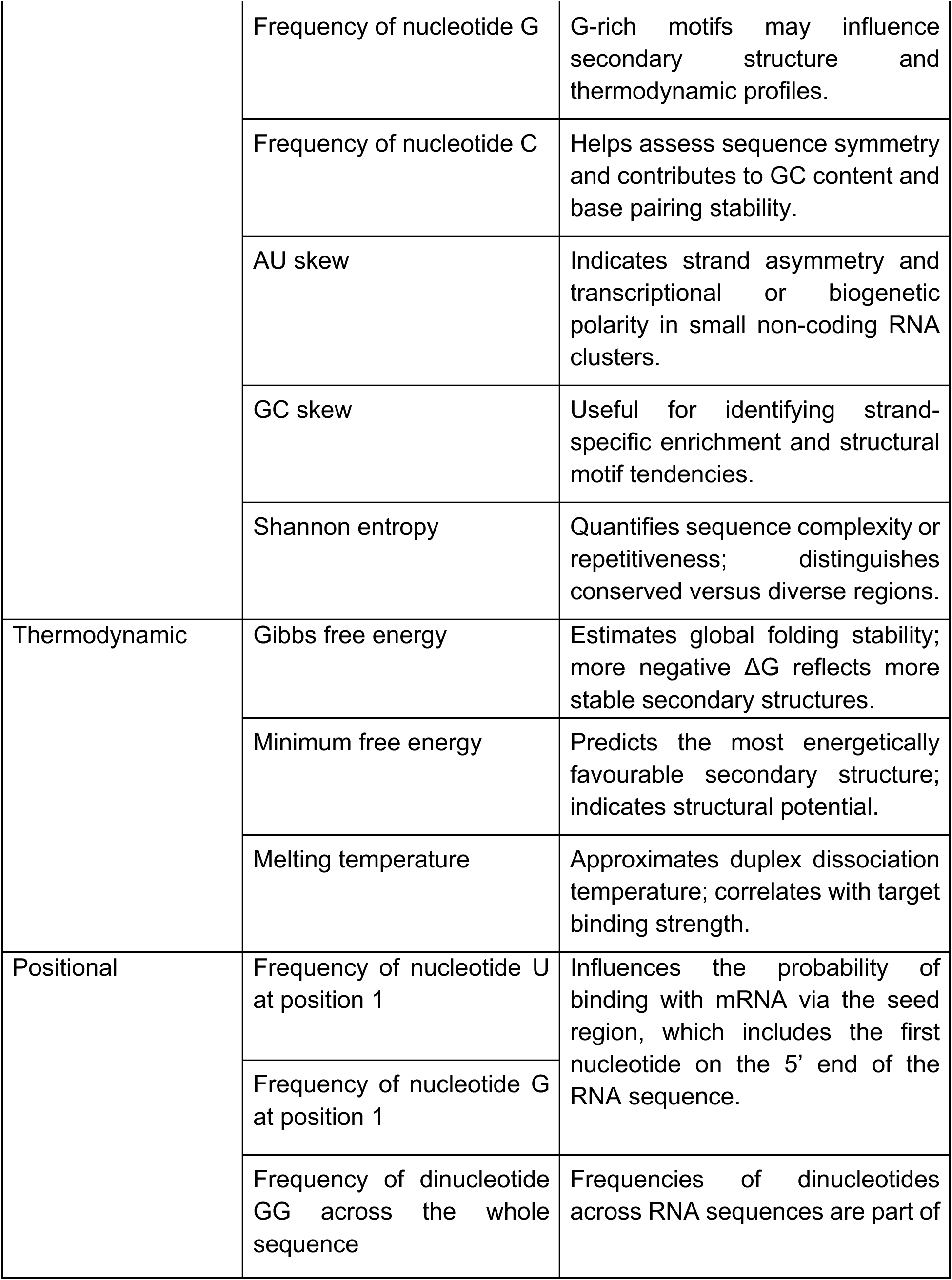
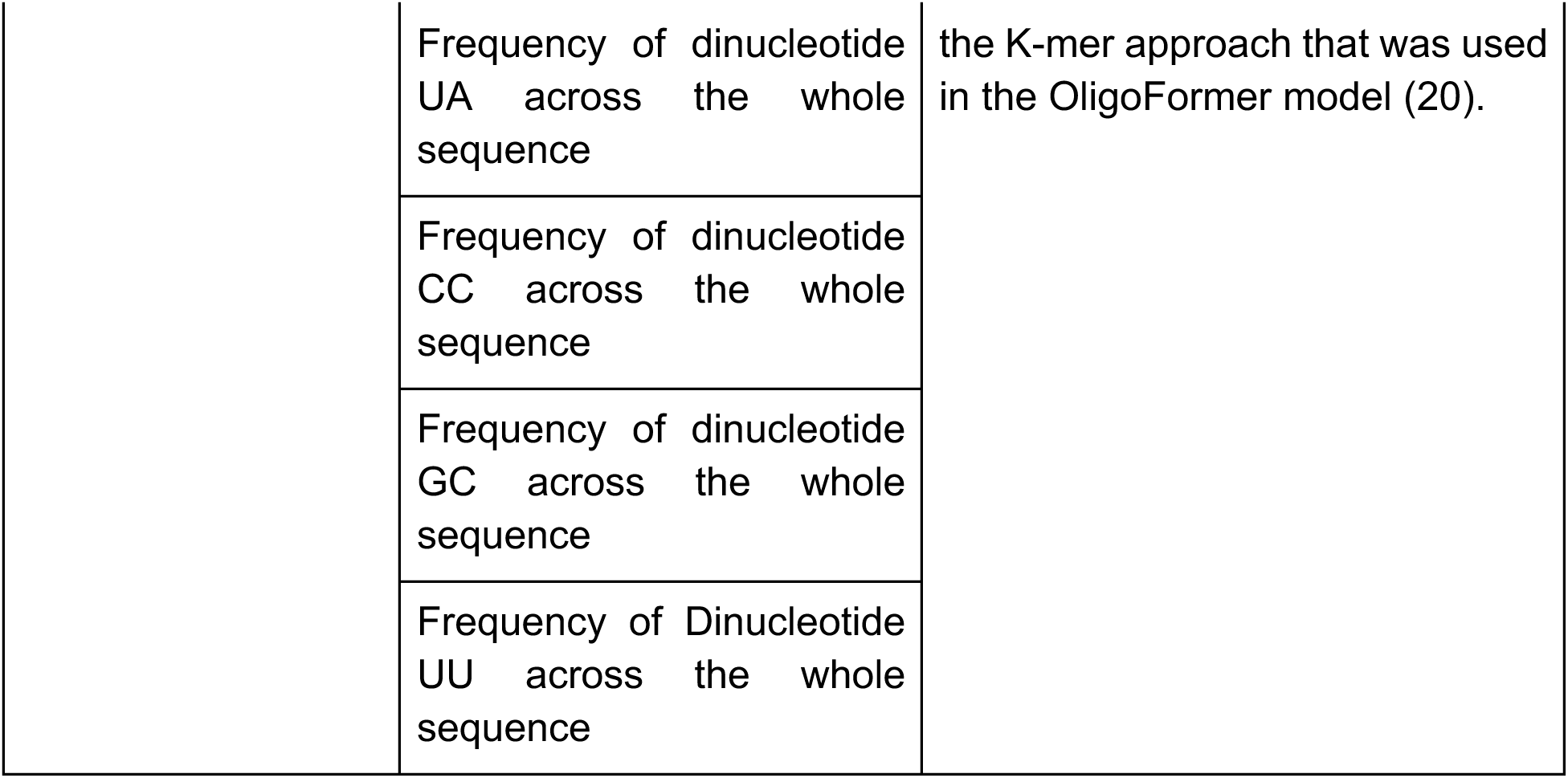
Details of the interpretable features used in the analysis.

| Category | Feature | Description |
| --- | --- | --- |
| Structural | GC content | Reflects duplex rigidity and contributes to structural stability and hybridisation potential. |
|  | Frequency of nucleotide A | Highlights base-specific biases relevant to miRNA seed regions and siRNA target pairing. |
|  | Frequency of nucleotide U | Whole-sequence uridine content; distinguishes uridine-rich synthetic guide strands from endogenous classes. |
|  | Frequency of nucleotide G | G-rich motifs may influence secondary structure and thermodynamic profiles. |
|  | Frequency of nucleotide C | Helps assess sequence symmetry and contributes to GC content and base pairing stability. |
|  | AU skew | Indicates strand asymmetry and transcriptional or biogenetic polarity in small non-coding RNA clusters. |
|  | GC skew | Useful for identifying strand-specific enrichment and structural motif tendencies. |
|  | Shannon entropy | Quantifies sequence complexity or repetitiveness; distinguishes conserved versus diverse regions. |
| Thermodynamic | Gibbs free energy | Estimates global folding stability; more negative $\Delta G$ reflects more stable secondary structures. |
|  | Minimum free energy | Predicts the most energetically favourable secondary structure; indicates structural potential. |
|  | Melting temperature | Approximates duplex dissociation temperature; correlates with target binding strength. |
| Positional | Frequency of nucleotide U at position 1 | Influences the probability of binding with mRNA via the seed region, which includes the first nucleotide on the 5' end of the RNA sequence. |
|  | Frequency of nucleotide G at position 1 |  |
|  | Frequency of dinucleotide GG across the whole sequence | Frequencies of dinucleotides across RNA sequences are part of |

|  |  |
| --- | --- |
| Frequency of dinucleotide UA across the whole sequence | the K-mer approach that was used in the OligoFormer model (20). |
| Frequency of dinucleotide CC across the whole sequence |  |
| Frequency of dinucleotide GC across the whole sequence |  |
| Frequency of Dinucleotide UU across the whole sequence |  |

Sequence composition features were derived from the nucleotide frequencies of adenine (A), uracil (U), guanine (G), and cytosine (C), normalised by sequence length. The GC content was computed as the proportion of nucleotides G and C relative to the total number of nucleotide bases, reflecting duplex rigidity and overall hybridisation potential. Strand asymmetry was captured using AU and GC skew, defined, respectively, as

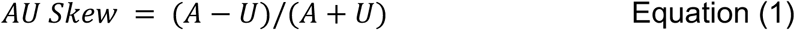

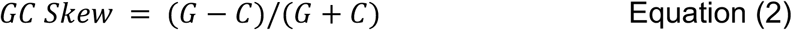

These ratios were calculated per sequence and reflect compositional bias across the sequence, useful for detecting asymmetry linked to transcriptional or processing directionality. To assess sequence complexity, Shannon entropy was computed over the nucleotide distribution using Equation 3,

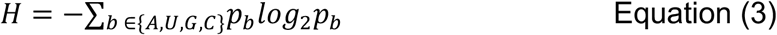

where *b* denotes a nucleotide base (A, U, G or C) and *p_b_* is computed based on the relative frequency of the base *b* in the sequence. Entropy values range from 0 bits (complete bias) to 2 bits (uniform distribution), capturing the variability or repetitiveness within each sequence. Biopython was used for sequence parsing. The resulting feature matrix was used for statistical analysis and comparative evaluation across RNA types. Other features utilised in previous models for predicting RNA efficiency were also considered. This includes seven features from Bai et al. (20) as they showed strong correlations with RNA-FM features (*r* ≥ 0.3 or *r* ≤ −0.3) and apply to all RNA types (Table 2). Correlation across all interpretable features was measured using Pearson correlation. A resulting heatmap was constructed, along with a network graph using the Kamada-Kawai algorithm (21).

To assess whether the distribution of structural motifs (stem versus bulge), as derived from predicted RNA secondary structure generated using ViennaRNA (RNAfold), varied across RNA types (siRNA, miRNA, piRNA), a chi-square test of independence was applied to the motif count data. Counts were arranged into a contingency table with motif types as rows and RNA types as columns. To understand the frequency of stem and bulge motifs across RNA types, both distribution of stem and bulge motifs per sequence and per nucleotide base were evaluated using the Kruskal-Wallis test with effect size measured by epsilon-squared. Dunn’s post-hoc test with Holm-Bonferroni correction was applied to measure pairwise significance.

### Computation of RNA-FM Features

Contextual embeddings for each RNA sequence were obtained using RNA-FM. The model consists of 12 transformer encoder layers and was trained using masked language modelling over a large corpus of RNA sequences to capture base-level and structural dependencies (12). For each input RNA sequence, RNA-FM generates a 640-dimensional representation for each nucleotide token. To standardise input representations, each small RNA sequence in the Data-23K dataset was passed through RNA-FM. Given that the model produces one embedding vector per nucleotide, average pooling was applied across the sequence length dimension to yield a fixed-length vector of size 640 for each sequence, regardless of original nucleotide count. This resulted in a final embedding matrix of shape 23,116 × 640 for each RNA type, with each row corresponding to a single RNA sequence embedding.

Dimensionality reduction of the embeddings was performed using t-distributed Stochastic Neighbour Embedding (t-SNE) applied to Data-23K. The t-SNE algorithm was configured with a perplexity of 30 and ran for 1000 iterations to reduce the 640-dimensional vectors to two-dimensional coordinates. Each point in the resulting 2D projection corresponds to an RNA sequence, and class labels (“siRNA”, “miRNA”, “piRNA”) were assigned to each point for plotting and analysis purposes. We plotted the t-SNE projection using scikit-learn. To examine how biologically interpretable features relate to the RNA-FM embedding space, the same t-SNE projection was coloured by interpretable features calculated per RNA type. Feature values were mapped onto each point in the 2D t-SNE space for visualisation purposes using consistent colour gradients. The analysis was repeated using Principal Component Analysis (PCA) using the first two components, given its deterministic nature. Silhouette score was employed to evaluate the clustering performance of t-SNE- and PCA-reduced RNA-FM embeddings using ground truth RNA type labels versus randomly assigned labels to investigate whether the clustering performances exceed random baselines.

To uncover RNA-FM’s utility in resolving subgroups within RNA types, we obtained miRNA family labels for Data-23K from miRBase (16,17). Clustering quality of RNA-FM embeddings across miRNA families was evaluated using average silhouette scores. Difference in clustering performance between ground truth miRNA family labels versus randomly assigned labels ground truth labels was evaluated with a two-sampled Kolmogorov-Smirnov test.

### Correlation Analysis

To analyse the relationship between all interpretable features and RNA-FM representations, we performed correlation analysis using Pearson correlation. For each interpretable feature, the 20 most positively and 20 most negatively correlated RNA-FM features were compiled, forming a total of 40 deep learning features. Correlations of each interpretable feature and the compiled RNA-FM features were then generated and clustered with hierarchical clustering (Euclidean method and Ward linkage). From the hierarchical clustering plot, we applied distance thresholds of 2.75 and 5 on the column and row dendrograms to identify clustering across interpretable features and deep learning features, respectively. Kernel Density Estimation (KDE) plots were generated to better visualise the correlation ranges of each feature. To further interrogate the correlation between deep learning-based features and interpretable features, a network graph was constructed with the two most positively and the two most negatively correlated RNA-FM features for each interpretable feature using the previously mentioned method.

All analyses were performed using Python, with random extraction performed using the built-in Random module, statistical analysis performed using pandas, SciPy and NumPy, and visualisation via Matplotlib, seaborn and NetworkX.

### RNA Type Classification and Prediction of Biological Features and siRNA Efficacy

The predictive power of RNA-FM embeddings from Data-23K were evaluated through RNA type classification and interpretable feature prediction. The dataset is split into 80% training and validation set and 20% held-out test set. With scikit-learn, a support vector machine was used for classification; whereas an ordinary least squares linear regression model was used for predicting continuous interpretable features. For binary features such as nucleotides U and G at position 1, a logistic regression model was implemented with 1000 iterations. In both tasks, 80% of the data were used for model training with 5-fold cross-validation, while the remaining 20% were held out as an independent test set for final performance evaluation. The metrics used to measure performance were accuracy, F1 score, precision and recall for classification, R^2^ score for linear regression, and ROC-AUC score for logistic regression. For siRNA efficacy prediction, linear regression models were trained using Huesken (13) and Takayuki (14) datasets, separately, with 10-fold cross validation. Due to the small sample size, no separate held-out test set was used. To prevent overfitting, we selected RNA-FM features that were highly correlated with interpretable features. Efficacy values were min-max scaled for each dataset.

### RNAExplorer web application

RNAExplorer is implemented as a web application using the Django framework, enabling modular development and interactive data visualisation within a browser-based environment. The backend is written in Python and manages data access, feature computation, and integration of reference embeddings, while the frontend provides an interactive interface for sequence input, visualisation, and tabular exploration of annotated small RNA datasets.

RNAExplorer supports both local and deployed execution modes. In the local installation, the Visualise Sequence feature allows user-provided RNA sequences to be processed on demand by computing RNA-FM embeddings, which are then projected into the reference embedding space of known small RNA sequences to enable real-time visualisation and comparison. In contrast, in the publicly deployed web version, the Visualise Sequence feature is restricted to user sequences that exactly match entries in the underlying reference database; such matched sequences are highlighted directly within the embedding plots.

RNAExplorer also allows users to find exact matches or fragment matches and retrieve 100 most similar sequences for a given sequence or fragment via the Query Sequence feature. The 100 most similar sequences are extracted from the reference database with the shortest Euclidean distance to the deep learning embeddings of a given sequence or fragment. Following the general approach of exponential similarity kernel (22), similarity is then represented by *exp(-d/σ)*, where *d* is the Euclidean distance between the deep learning embeddings of the given sequence or fragment and the reference sequence, and *σ* is a fixed scaling parameter (*σ* = 1).

## RESULTS

### Small RNA classes exhibit distinct structural and thermodynamic properties

We first sought to systematically characterise and compare the compositional, positional, structural and thermodynamic properties of siRNAs, miRNAs, and piRNAs. This analysis was designed to identify shared and RNA-class-specific patterns, and to provide an interpretable reference for subsequent assessments of how these properties are represented within RNA-FM embeddings. This analysis identified both shared relationships and distinct class-specific feature distributions (Fig. 2). ΔG, MFE and T_m_ distributions demonstrated significant differences among RNA types (Fig. 2a–c, Table 3). The most negative ΔG and MFE values were exhibited by piRNAs, indicating greater thermodynamic stability, followed by miRNAs and then siRNAs. Consistent with this, piRNAs also have the highest melting temperature profile, followed by miRNAs and siRNAs, reflecting a more stable structure. Shannon entropy values, reflecting sequence complexity, were moderately elevated in piRNAs compared to miRNAs and siRNAs, suggesting greater sequence variability (Fig. 2d, Table 3).

**Figure 2.**
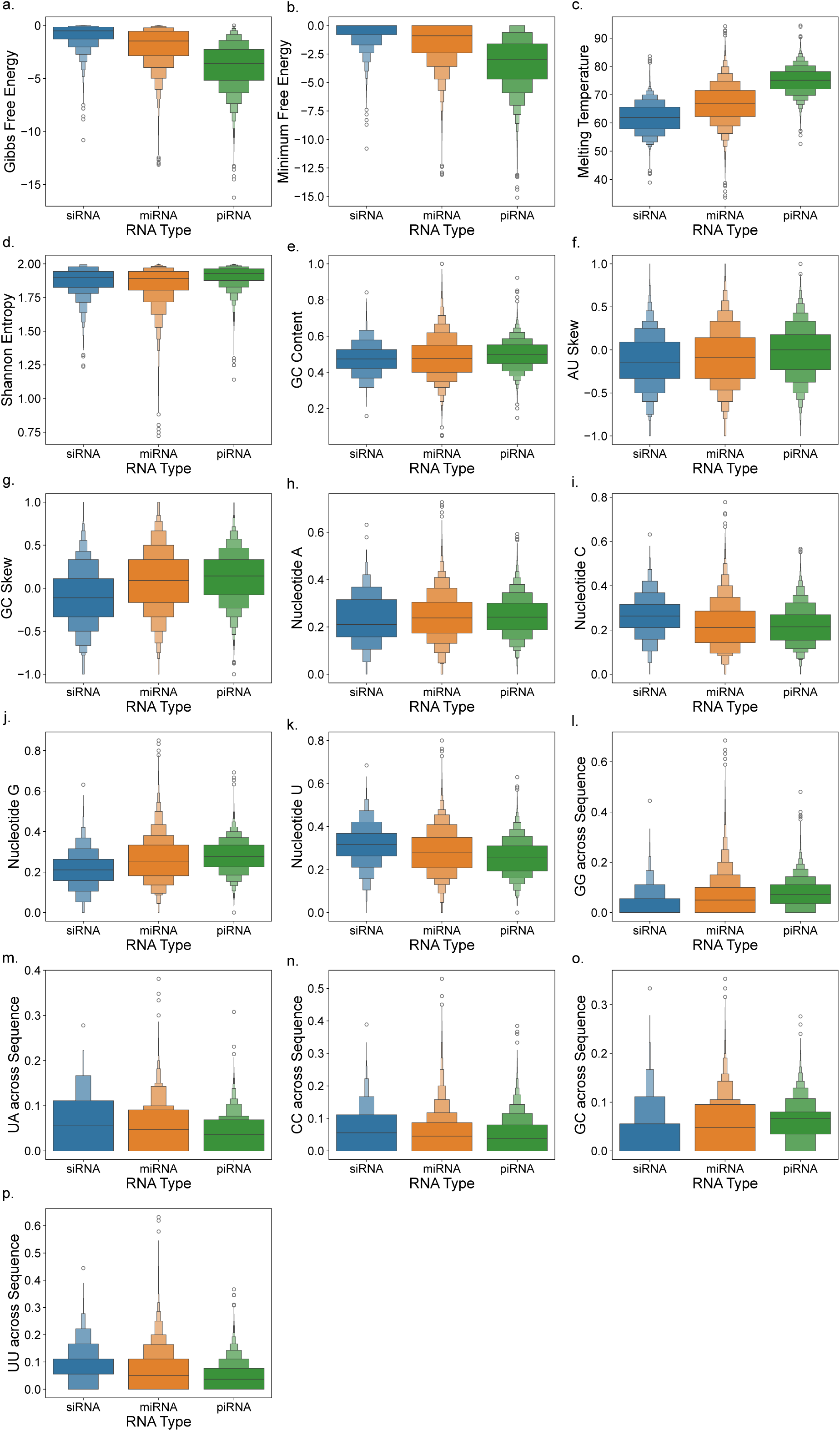
Density distribution plots for all extracted interpretable features, namely (a) Gibbs Free Energy, (b) Minimum Free Energy, (c) Melting temperature, (d) Shannon Entropy, (e) GC content, (f) AU Skew, (g) GC Skew, (h–k) nucleotides A, C, G, U, (I–p) dinucleotides GG, UA, CC, GC, UU across sequence for siRNAs (blue), miRNAs (orange) and piRNAs (green), respectively. All features are significantly different across all three RNA types as assessed via a two-sample Kolmogorov-Smirnov test (p <0.00001).

**Table 3.** Summary statistics of key features for siRNA, miRNA, and piRNA across Data-23K.

| Feature | siRNA (Mean ± Standard Deviation) | miRNA (Mean ± Standard Deviation) | piRNA (Mean ± Standard Deviation) |
| --- | --- | --- | --- |
| Gibbs Free Energy, $\Delta G$ (kcal/mol) | -0.88 ± 1.01 | -1.92 ± 1.74 | -3.83 ± 2.12 |
| Minimum Free Energy, MFE (kcal/mol) | -0.57 ± 1.00 | -1.50 ± 1.77 | -3.24 ± 2.21 |
| Melting Temperature, $T_m$ (°C) | 61.72 ± 5.29 | 66.89 ± 6.95 | 75.06 ± 4.55 |
| Shannon Entropy | 1.88 ± 0.10 | 1.85 ± 0.13 | 1.91 ± 0.07 |
| GC Content (%) | 46.0 ± 9.0 | 48.0 ± 12.0 | 50.0 ± 8.0 |
| AU Skew | -0.13 ± 0.32 | -0.08 ± 0.34 | -0.02 ± 0.30 |
| GC Skew | -0.12 ± 0.35 | 0.09 ± 0.38 | 0.13 ± 0.31 |
| Sequence length (nt) | 19 | ~22 | ~29 |

Next, we sought to analyse sequence composition across different RNA types. Both higher AU and GC skew values are observed in piRNAs compared to miRNAs and siRNAs (Fig. 2f, g). Mononucleotide distributions also revealed RNA-class-specific biases. piRNAs were enriched in nucleotides A and G and depleted in nucleotide C relative to miRNAs and siRNAs (Fig. 2h–j). Consistent with their G enrichment, piRNAs also showed higher frequencies of GG and GC dinucleotides (Fig. 2j, l, o), reflecting their overall purine-rich sequences. By contrast, the U-rich composition of siRNAs was reflected in their greater frequencies of UA and UU dinucleotides, despite their comparatively low A content (Fig. 2h, k, m, p). Overall, the global feature distributions demonstrate distinct thermodynamic and compositional profiles among the three small-RNA classes. piRNAs were characterised by greater thermodynamic stability and higher GC content, whereas siRNAs were comparatively U-rich and less thermodynamically stable, with miRNAs generally showing intermediate profiles.

### Correlation Patterns Among Interpretable Features

Correlation analysis between interpretable features in Data-23K revealed a group of related features (Fig. 3a, b). Gibbs free energy and minimum free energy were significantly correlated (*r* ≈ 0.99, *p* < 0.001), indicating close alignment between total and minimum folding energy values. Additionally, melting temperature and GC content are positively correlated (*r* ≈ 0.78, *p* < 0.001). Melting temperature and GC content showed strong negative correlations with Gibbs free energy and minimum free energy (For T_m_, ΔG: *r* ≈ −0.69, MFE: *r* ≈ −0.65, while for GC content, ΔG: *r* ≈ −0.44, MFE: *r* ≈ −0.43; all *p* < 0.001), while UA motifs across the sequence showed an opposite trend (ΔG: *r* ≈ 0.27, MFE: *r* ≈ 0.25, *p* < 0.01). Sequence length was positively correlated with melting temperature (*r* ≈ 0.70, *p* < 0.001) and negatively correlated with minimum free energy (*r* ≈ −0.56, *p* < 0.001), consistent with longer sequences forming a more stable secondary structure. Correlation patterns in the human-only Data-3K largely mirrored those in the Data-23K (Supplementary Fig. S2).

**Figure 3.**
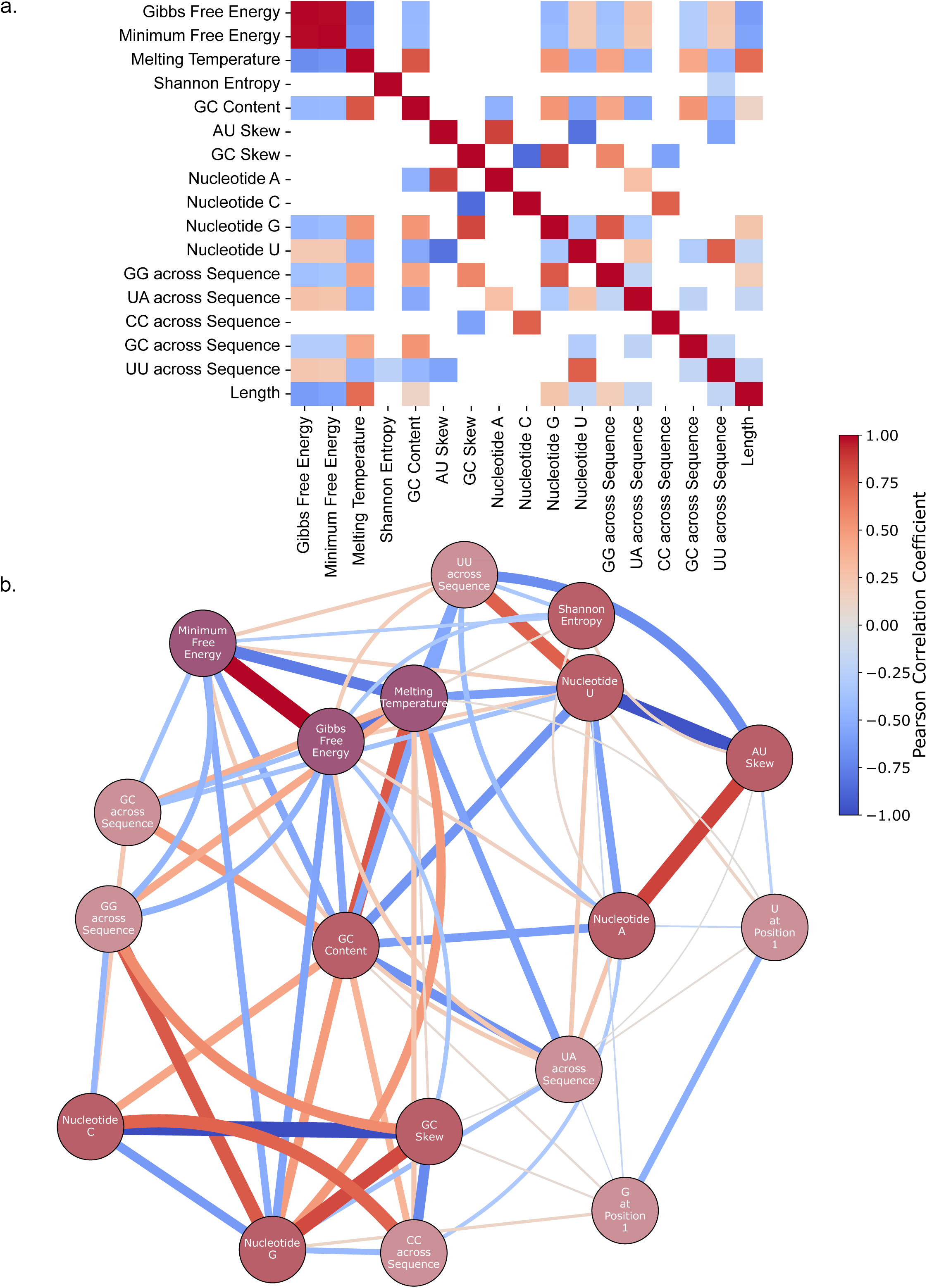
Feature correlation analysis across Data-23K. (a) Correlation heatmap illustrating pairwise Pearson correlation coefficients among extracted features, with non-significant correlations (*p* ≥ 0.05) left blank. (b) Network graph showcasing the three most positively and the three most negatively correlated features of each interpretable feature for all RNA sequences. All interpretable features are categorised into structural (brick red), positional (light pink), and thermodynamic (dark purple) features. Edge colour represents the Pearson correlation coefficient as shown by the colour bar, whereas edge weight represents the degree of correlation.

### Structural Motif Distribution

We observed that different RNA types showed generally similar correlation patterns in nucleotide frequencies (Fig. 4a–c). Quantitative comparison of structural motifs formed within the sequence revealed that stem motifs appear at a substantially higher frequency than bulge motifs across all RNA types (Fig. 4d). Notably, siRNAs showed the highest stem-to-bulge motif ratio while piRNAs showed the lowest stem-to-bulge motif ratio (chi-square *p*-value < 0.0001). Distribution of stem and bulge motif counts per sequence revealed that piRNA has more stem and bulge motifs in a sequence than miRNA and siRNA, with Kruskal-Wallis test showing significant differences (Stem motifs: *H*(2) = 20011.80, *p* < 0.0001, *ε^2^* = 0.2886; Bulge motifs: *H*(2) = 3783.80, *p* < 0.0001, *ε^2^* = 0.0546) (Fig. 4e, g). Dunn’s post-hoc test with Holm-Bonferroni correction confirmed significance across all three RNA types (*p* < 0.0001 for all three pairs) (Fig. 4e, g). To account for sequence length effect, the distributions of stem and bulge motif fraction per nucleotide base confirmed piRNA’s inherent tendency to form more complex self-folding structures, compared to miRNA and siRNA (Fig. 4f, h). Both stem and bulge motifs show significant difference using Kruskal-Wallis test (Stem motifs: *H*(2) = 7380.61, *p* < 0.0001, *ε^2^* = 0.1064; Bulge motifs: *H*(2) = 3390.39, *p* < 0.0001, *ε^2^* = 0.0489), with pairwise significance across all RNA types observed with Dunn’s post-hoc test (*p* < 0.0001 for all three pairs) Across three RNA types, piRNA sequences have higher structural complexity, as evidenced by higher stem motifs; whereas synthetic siRNA in our dataset has minimal self-folding and tends to retain less complex structures.

**Figure 4.**
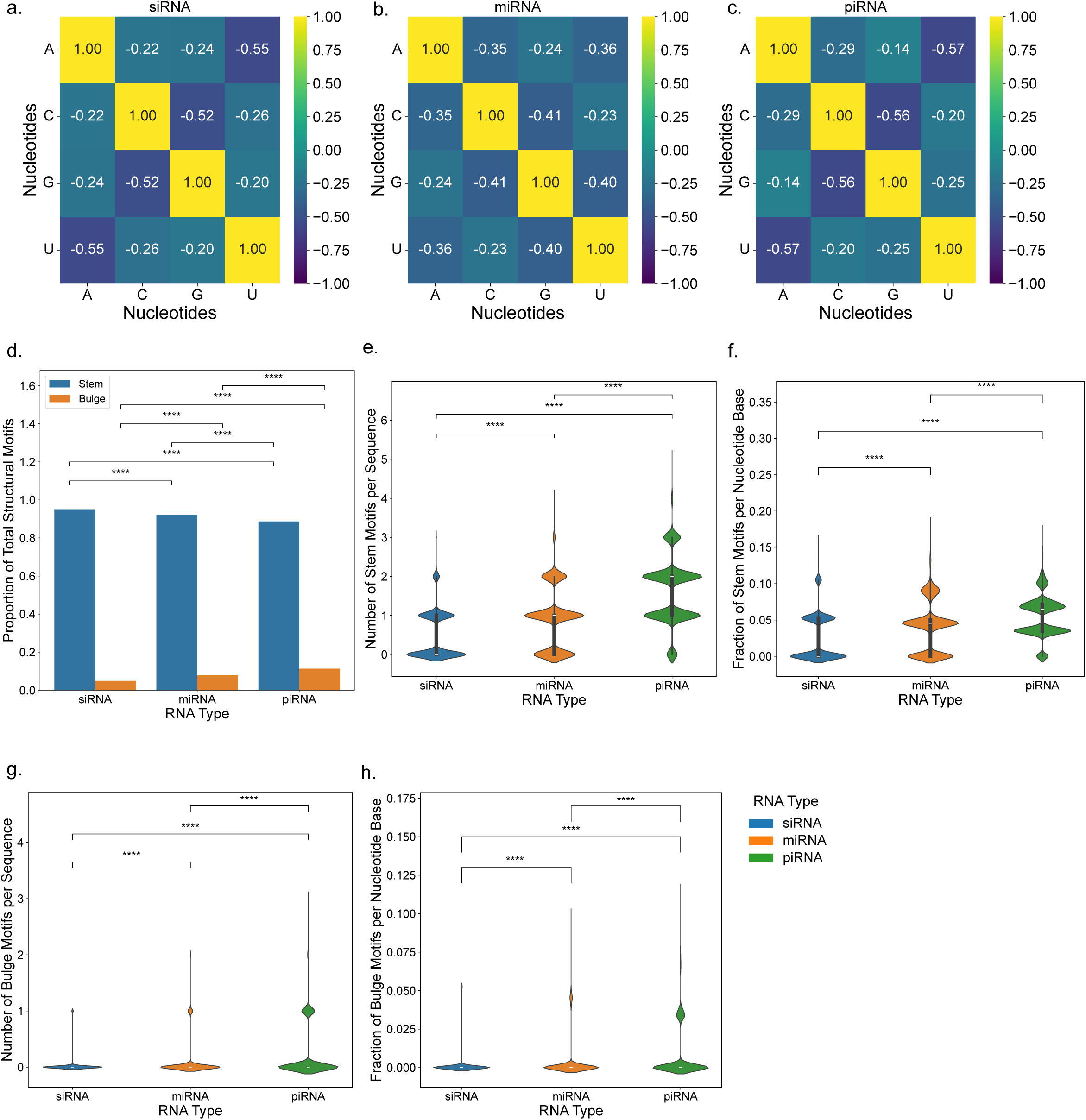
Nucleotide correlation matrices and structural motifs comparison for different small RNA types. (a–c) Pearson correlation metrics showing pairwise relationships among nucleotide proportions (A, C, G, and U) for siRNA, miRNA, and piRNA, respectively. (d) Proportion of stem (blue) and bulge (orange) motifs relative to total number of structural motifs, summed within each RNA type, with pairwise significance tested with chi-squared test of independence. (e) Distribution of stem motifs count per sequence across RNA types, with significance tested with a Kruskal-Wallis test followed by Dunn’s post-hoc test (*H*(2) = 20011.80, *p* < 0.0001, *ε^2^* = 0.2886). (f) Distribution of fraction of stem motifs per nucleotide base across RNA types, with significance tested with a Kruskal-Wallis test followed by Dunn’s post-hoc (*H*(2) = 7380.61, *p* < 0.0001, *ε^2^* = 0.1064). (g) Distribution of bulge motifs count per sequence across RNA types, with significance tested with a Kruskal-Wallis test followed by Dunn’s post-hoc test (*H*(2) = 3783.80, *p* < 0.0001, *ε^2^* = 0.0546). (h) Distribution of fraction of bulge motifs per nucleotide base across RNA types, with significance tested with a Kruskal-Wallis test followed by Dunn’s post-hoc (*H*(2) = 3390.39, *p* < 0.0001, *ε^2^* = 0.0489). ****, *p* < 0.0001.

### RNA-FM features reveal class-specific patterns across small RNAs

Having established the shared and class-specific patterns of interpretable properties across siRNAs, miRNAs and piRNAs, we next investigated whether information associated with these properties was represented within RNA-FM embeddings generated from sequence alone. During pretraining, RNA-FM learns to predict masked nucleotides from their surrounding sequence context, enabling the model to capture recurring sequence patterns and dependencies. The resulting output is a high-dimensional vector representation of each RNA sequence, referred to as an embedding. In representation learning, information associated with an RNA property is considered encoded when it is represented within, and can be predicted from, the learned embedding.

To visualise the RNA-FM embeddings for 69,348 small RNA sequences, we used t-distributed Stochastic Neighbour Embedding (t-SNE) which projects the 640-dimensional representations into two dimensions while placing sequences with locally similar embeddings near one another. The resulting local groupings indicate similarity in the sequence information captured by RNA-FM. The resulting distribution shows a clear spatial separation between piRNAs and the other two RNA types (Fig. 5d), where piRNAs formed a distinct and densely populated cluster (Fig. 5c). In contrast, siRNAs and miRNAs occupied overlapping but partially distinguishable regions on the right side of the projection (Fig. 5a, b). The broadest region appeared to be occupied by miRNAs, with siRNAs showing intermediate spread and piRNAs occupying the most compact region. The overall structure of the t-SNE projection shows a continuous transition between these groups while retaining RNA-type-specific patterns represented within the RNA-FM embeddings.

**Figure 5.**
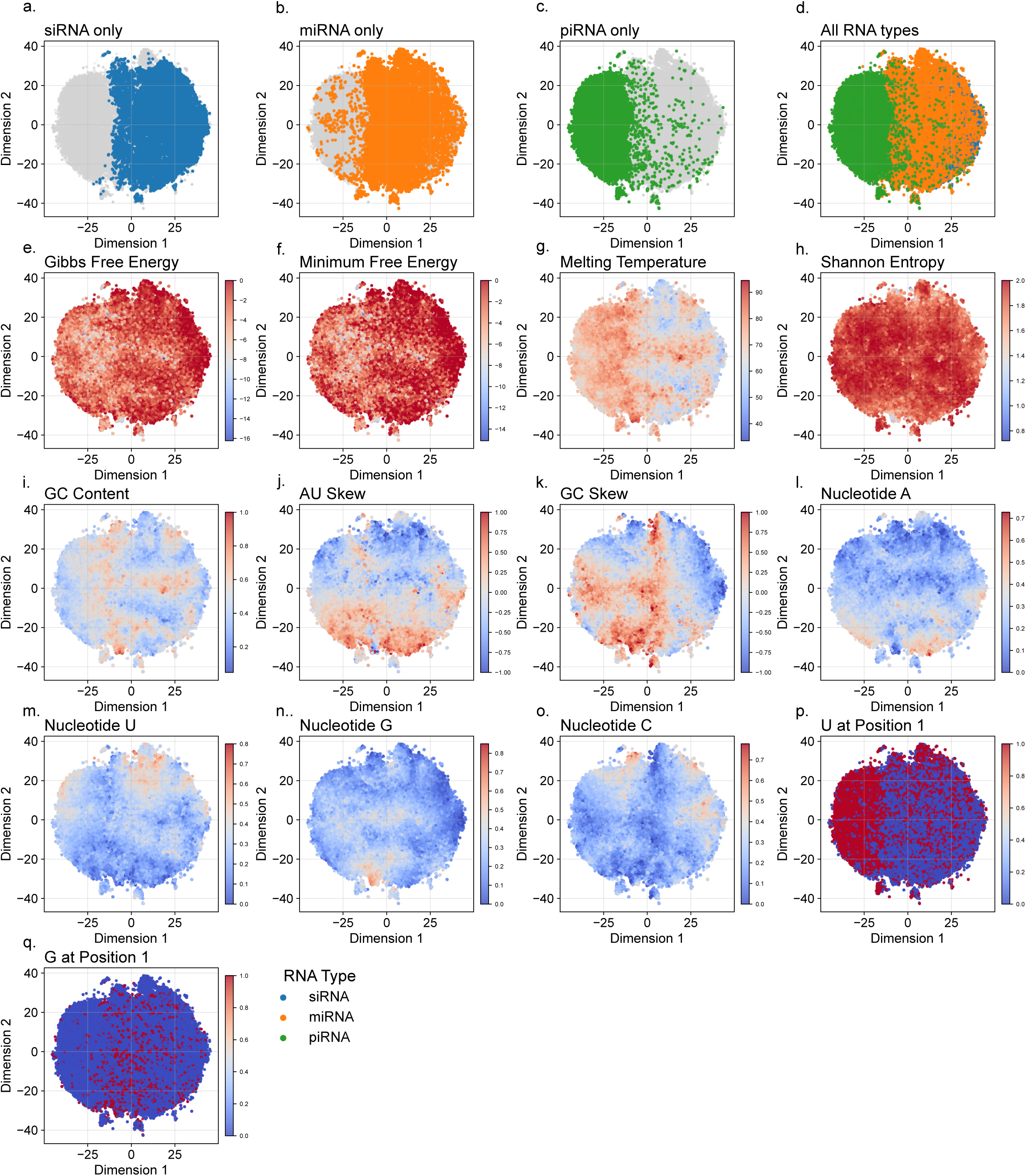
t-SNE visualisation of RNA-FM embeddings across all RNA types and feature annotations. All panels show the same two-dimensional t-SNE projection of RNA embeddings. Sequences are highlighted by RNA class in (a) siRNA (blue), (b) miRNA (orange) and (c) piRNA (green), and all three classes are shown together in (d). In (e–q), sequences are coloured according to the values of each interpretable feature with blue hues indicating lower feature values and red hues indicating higher feature values.

As different RNA types varied in their length, we assessed whether sequence length accounted for the organisation of the embedding space (Supplementary Fig. S3). Across all RNA sequences, length was strongly correlated with t-SNE dimension 1 (*r* ≈ −0.82, *p* < 0.001; Supplementary Fig. S3e). However, this association was substantially weaker within individual RNA types (miRNA: *r* ≈ −0.21; piRNA: *r* ≈ −0.18; both *p* < 0.001; Supplementary Fig. S3f–h), suggesting that the strong global correlation largely reflected differences in length distributions among RNA classes. Sequence length was not correlated with t-SNE dimension 2 (*r* ≈ −0.02; Supplementary Fig. S3i–l).

We further examined length-matched subsets to distinguish the effect of sequence length from RNA-type-specific characteristics. Specifically, we compared siRNAs and miRNAs of 19 nt, and piRNAs and miRNAs of 26–32 nt (Supplementary Fig. S3m, n). Despite having matched lengths, sequences from different RNA types retained distinct spatial distributions. Moreover, sequences of the same length could occur in different regions of the embedding: 19-nt sequences were present within the piRNA-associated region, whereas 26–32-nt sequences were also found outside that region. Some piRNAs additionally overlapped the siRNA- and miRNA-associated regions (Supplementary Fig. S3d). Together, these analyses indicate that sequence length contributes to the global organisation of the embedding but does not fully account for the separation among RNA types. A similar organisation was observed in the human-only Data-3K dataset, including a compact, left-shifted piRNA cluster and overlapping miRNA and siRNA regions.

To examine how interpretable features align with the RNA-FM embedding structure, the same t-SNE projection was coloured by feature values independently calculated for each RNA sequence (Fig. 5e–q). Gibbs free energy and minimum free energy showed similar spatial patterns, with lower values concentrated in the left region of the embedding corresponding to the piRNA region and higher values toward the right, occupied by siRNAs and miRNAs (Fig. 5e, f). In contrast, the melting temperature showed an inverse trend, with higher values presented in the piRNA region and lower values in the siRNA and miRNA areas (Fig. 5g). Nucleotide U at position 1 showed a strong and well-defined enrichment in the left half of the map (Fig. 5p). This left-side concentration aligns with the piRNA cluster, suggesting a high frequency of 5′-uridine in piRNAs. The right half of the embedding, which corresponds to siRNAs and miRNAs, displays noticeably lower levels of U at the first position. These differences demonstrated how RNA-FM features capture biological characteristics of small RNA sequences and that the separation of piRNA sequences in the latent space was likely due to differences in both composition and thermodynamic features.

Shannon entropy was largely uniform across the embeddings with only minor local variation (Fig. 5h). GC content was generally low, with small hotspots but no strong region-specific accumulation (Fig. 5i). In contrast, AU and GC skew showed clear spatial structure: AU skew was highest in the bottom region of the landscape consistent with relative enrichment for adenine, and most negative in the upper-left region, consistent with relative enrichment for uridine (Fig. 5j). GC skew tended to be higher in the bottom and central regions (G-rich) and generally lower in the left region (C-rich) (Fig. 5k).

Nucleotide maps mirrored these patterns where nucleotide A was enriched in the lower right and depleted in upper regions (Fig. 5l); nucleotide U was enriched in the upper centre and left regions and reduced in the bottom half (Fig. 5m); nucleotide G showed a small, localised hotspot in the bottom central region with low values elsewhere (Fig. 5n); nucleotide C was modestly enriched in the top left and top right, and otherwise low (Fig. 5o). For positional nucleotides, nucleotide U at position 1 showed a complementary enrichment pattern, whereas nucleotide G at position 1 was sparse and broadly dispersed with no dominant focus (Fig. 5p, q). Together, the t-SNE projections of RNA-FM embeddings coloured by interpretable features confirmed that RNA-FM features captured many of these features in a non-linear manner.

### Associations between RNA-FM dimensions and interpretable RNA features

We then investigated the biological representation of RNA-FM embeddings by correlating interpretable features and individual deep learning features. Hierarchical clustering of the correlation values between interpretable and RNA-FM features across all RNA types provided direct interpretations of some of these features (Fig. 6). To better understand these correlation patterns, we defined two cluster types by clustering interpretable and deep learning features using distance thresholds on the column and row dendrograms: interpretable Cluster (iCls) and deep learning Cluster (dCls). The column and row dendrograms revealed 7 distinct iCls and 21 distinct dCls. Except for minimum free energy and Gibbs free energy, distinct correlation patterns were observed between different deep learning features and interpretable features, demonstrating how RNA-FM can capture distinct structural and chemical features. For example, the correlation values for nucleotide U were highly similar to UU across the sequence, with slight variation where the latter had lower correlation values (iCls1). Interestingly, these correlation values were distinct from correlation values with nucleotide U at position 1 (iCls4), showcasing that RNA-FM distinguished positional nucleotides from sequence-wise nucleotide information. Similarly, correlation patterns of GC content and dinucleotide GC across the sequence (iCls5) were highly similar except for deep features in dCls11, 15, 17, and 20. Although Shannon entropy did not show a clear trend in t-SNE landscape, it was grouped in iCls4 along with positional nucleotide features and showed moderate correlation with several single deep learning-based features. Thermodynamic features also showed similar correlation patterns to compositional features, such as melting temperature with GC content and dinucleotide GC across sequence (iCls5), and Gibbs free energy and minimum free energy with dinucleotide UA across sequence (iCls2), providing important considerations of how these features could interact. These analyses provide insights on associated interpretable features in small RNA and support the interpretation of deep learning features, which could provide scientists with a wide range of accessible features.

**Figure 6.**
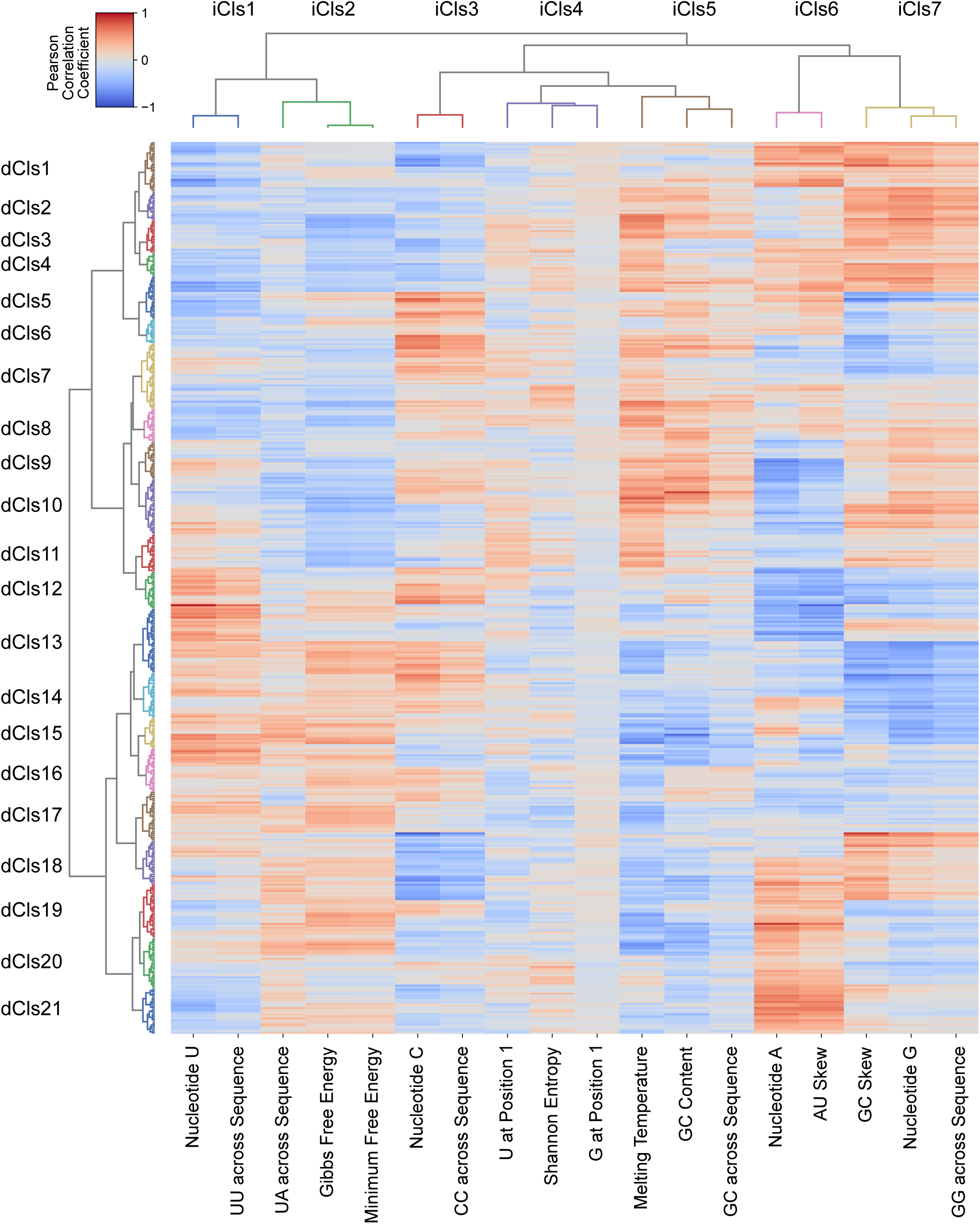
Heatmap of Pearson correlations between interpretable features and RNA-FM features with hierarchical clustering for the All Types data from Data-23K. Each cluster is coloured based on distance thresholds. iCls: interpretable feature clusters; dCls: deep learning feature clusters.

To assess whether the relationships between RNA-FM embeddings and interpretable features differ across RNA types, Pearson correlation patterns were plotted separately for siRNA, miRNA, and piRNA (Supplementary Fig. S4 and S5). Most correlation values were highly similar across RNA types, except for Shannon entropy and nucleotides G and U at position 1, which showed generally higher correlations with deep learning features in miRNA and siRNA sequences, and piRNA sequences, respectively. These patterns from Figure 6 and Supplementary Fig. S4 were consistent when human-only sequences as Data-3K were considered, except that nucleotide G at position 1 now forms iCls7 with dinucleotide GC across sequence (Supplementary Fig. S6 and S7). These observations suggest complex interactions between some of the interpretable features and that deep learning can capture higher-level sequence features associated with each RNA type.

To visualise the distribution of correlation values between interpretable features and deep features across different RNA types, we applied Kernel Density Estimation (KDE) on Data-23K (Fig. 7). Generally, correlation values had similar distributions across different RNA types. Structural features from nucleotides A, U, G, and C to GC content predominantly had wide to moderate peaks depicting high to medium correlations (−0.8–0.8). On the other hand, Shannon entropy distribution had a narrow peak, suggesting low correlations (−0.3–0.3) with piRNA’s Shannon entropy having the narrowest peak. Similarly, nucleotides U and G at position 1, with narrow peaks overall, especially for siRNA and miRNA sequences, suggest low correlation with RNA-FM embeddings. Gibbs free energy and minimum free energy also had lower correlation values for the individual types, but higher correlation values were observed when the three RNA types were combined. The melting temperature had more moderate peaks along with a stronger correlation compared to the two free energy variables. Similar patterns were observed in Data-3K (Supplementary Fig. S8). These correlation ranges suggested that RNA-FM’s can capture biological information associated with mononucleotide positional features, thermodynamic features, dinucleotide positional features and structural features. Shannon entropy and melting temperature were both outliers, with the former behaving similarly to mononucleotide positional features and the latter identical to structural features such as GC content.

**Figure 7.**
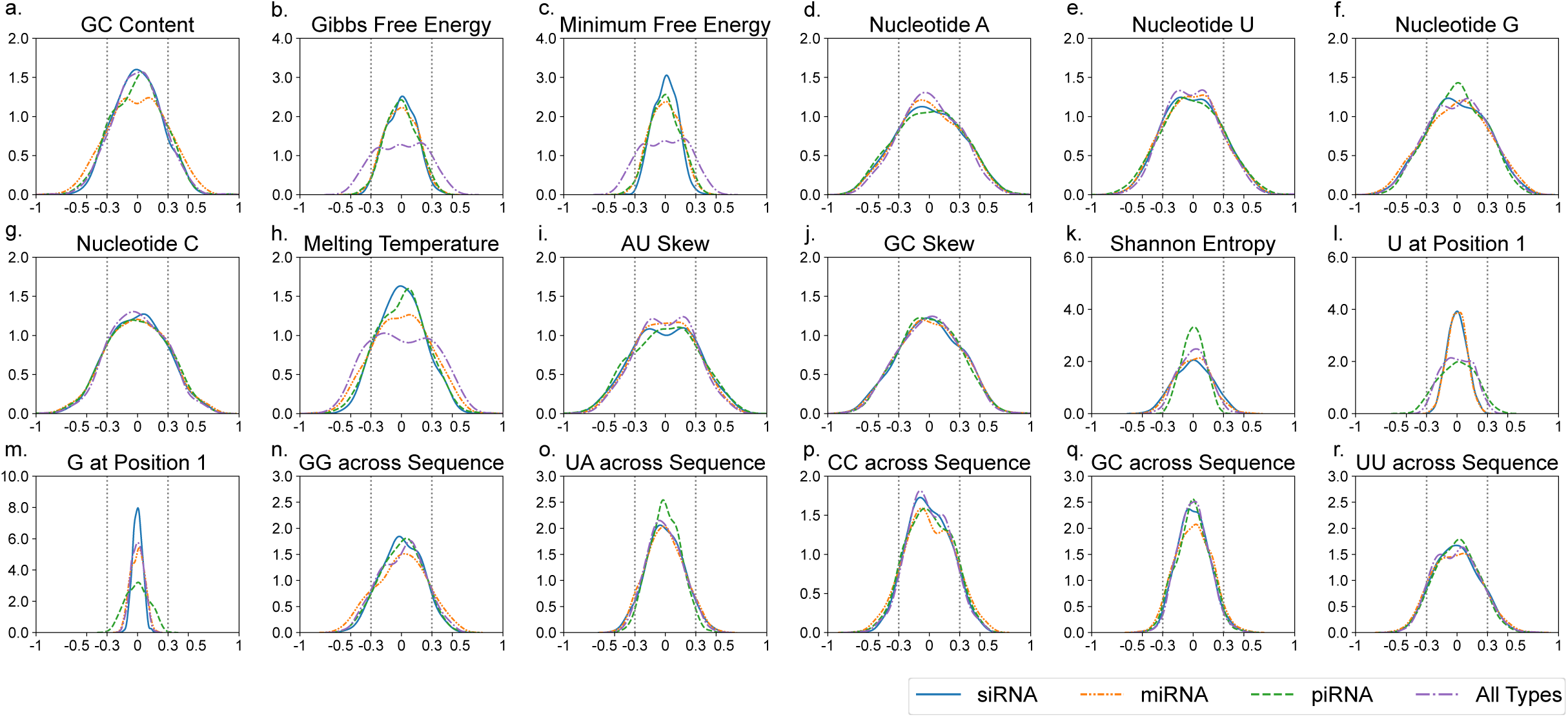
Kernel Density Estimation of Pearson correlation values (x-axis) of each interpretable feature for siRNAs (blue), miRNAs (orange), piRNAs (green) and All Types (purple). Dotted vertical lines indicate correlation values of −0.3 and 0.3.

### Correlation mapping links RNA-FM dimensions to interpretable properties

Next, we sought to determine specific RNA-FM features that can provide a proxy for various interpretable features. To achieve this, we investigated RNA-FM features that showed the strongest correlations with interpretable features across different RNA types. For each interpretable feature, we selected the two most positively and the two most negatively correlated deep learning features for All Types, siRNA, miRNA, and piRNA only (Fig. 8). Interpretable features were linked in the graph through their correlations with common deep learning features. Consequently, certain relationships previously presented in Figures 3 and 6 were recapitulated. For instance, GC content and melting temperature formed a tightly connected pair through their shared association with features 148 and 90 across siRNA, miRNA, and piRNA. Likewise, the negative correlation between GC content, Gibbs free energy, and minimum free energy for the three RNA types can be inferred from their shared associations with features 148, 90 and 116. In contrast, Shannon entropy appeared disconnected from other interpretable features due to distinct top correlated deep learning features. As for nucleotides U and G at position 1, they shared three deep learning features, suggesting that these features encode positional information, whereas the remaining 17 correlated features likely capture nucleotide-specific signals.

**Figure 8.**
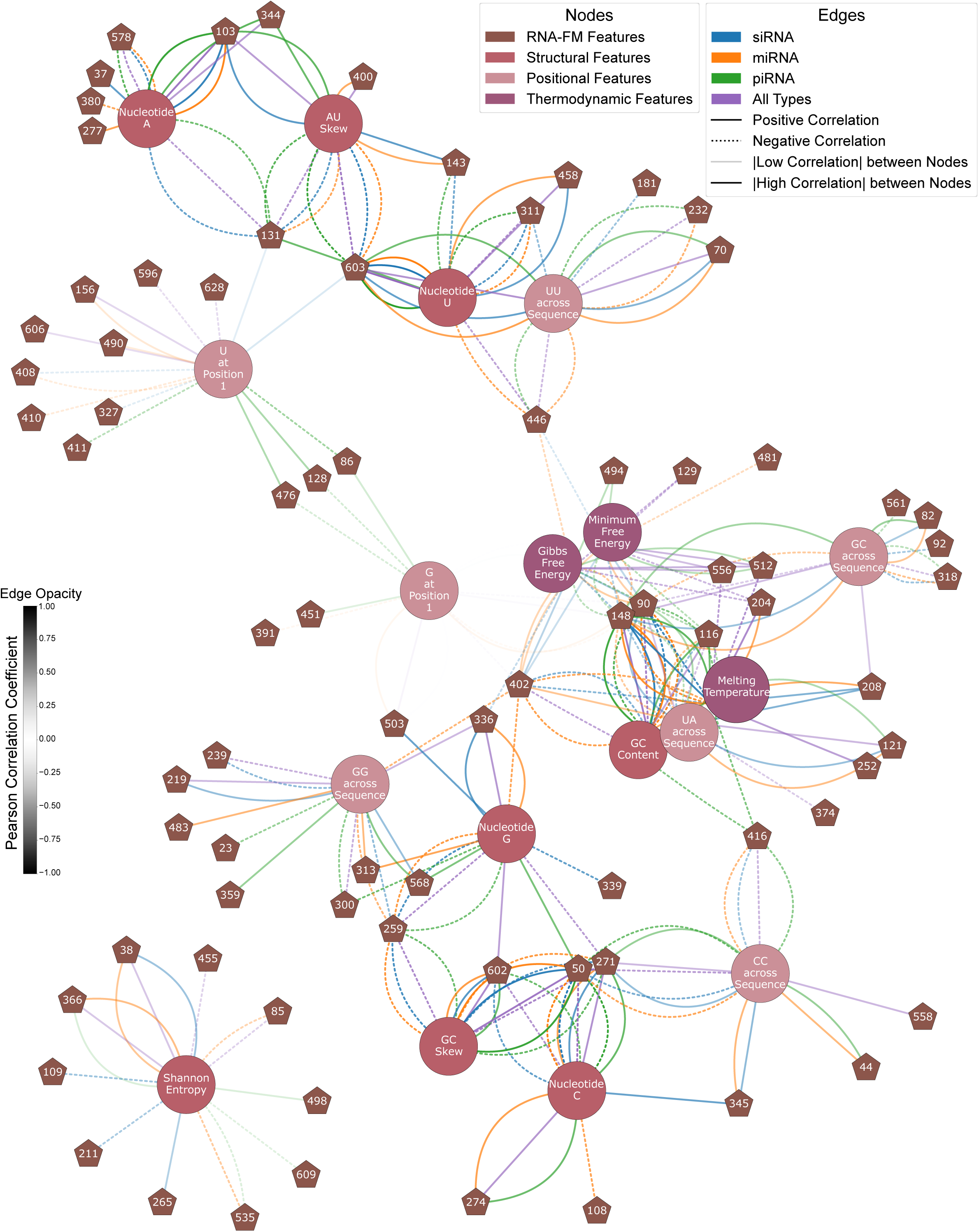
RNA-FM features can distinguish RNA-specific properties. Network graph showing the two most positively and the two most negatively correlated RNA-FM features (brown) to each interpretable feature. All interpretable features are categorised into structural (brick red), positional (light pink) and thermodynamic (dark purple) features. All edges are coloured according to RNA type: siRNA (blue), miRNA (orange), piRNA (green), and All Types (purple). Edge opacity represents the degree of Pearson correlation as shown by the colour bar. Solid edges represent positive correlation, whereas dotted edges represent negative correlation.

It is observed that some deep learning representations correlate strongly with specific interpretable features both within individual RNA types and across the combined dataset (“All Types”). Examples include feature 259, which correlated with Nucleotide G and GC skew, feature 103, which correlated with Nucleotide A, and feature 148, which correlated with GC content and GC across sequence. Figure 8 also shows that some of the deep learning features are strongly correlated with certain RNA types. An example is feature 402 that correlates with seven interpretable features in miRNA but only five interpretable features in siRNA. Although feature 402 appears to not strongly correlate with any interpretable features in piRNA, closer inspection reveals that it still ranks within the ten most positively or negatively correlated features with seven interpretable features for piRNA. We can also pinpoint which biological characteristics certain deep learning features are most correlated with, such as the correlation between feature 603 and uridine-related features. Interestingly, while feature 50 correlates with both guanine- and cytosine-related features, it does not highly correlate with GC content. These results further support that RNA-FM-derived features can distinguish RNA-type-specific features and offer important insights into explaining their success in various predictive tasks (12, 20, 23).

### RNA-FM representations support classification and feature prediction

We conducted several prediction tasks to evaluate the predictive power of deep learning embeddings compared to interpretable features. An RNA type classification task was conducted to confirm that the embeddings contain useful RNA-type-specific information (Supplementary Fig. S9, Supplementary Table S1). On the held-out test set, RNA-FM embeddings improved the F1-score from 0.935 to 0.969 compared with interpretable features only, with consistent improvements across all performance metrics. The largest improvement was observed for miRNA classification, where the F1-score increased from 0.898 using interpretable features to 0.953 using RNA-FM embeddings (Supplementary Fig. S9b). These results further support that RNA-FM embeddings encode meaningful type-specific biological characteristics.

To further confirm that the separation between different small RNA types observed in the t-SNE space is reproducible, we measured the clustering coefficient for each RNA type in the t-SNE and the Principal Component Analysis (PCA) reduced latent space. Clustering performance of t-SNE-reduced RNA-FM embeddings using ground truth RNA type labels was better compared to randomly assigned labels, as shown by the higher silhouette score (Supplementary Table S2), confirming that RNA-FM embeddings capture type-specific biological information, as observed in the t-SNE space (Fig. 5). Similar results were observed for PCA-reduced RNA-FM embeddings, supporting the robustness of t-SNE projection (Supplementary Fig. S10 and Supplementary Table S3).

We also evaluated the utility of RNA representations in resolving miRNA subgroups by investigating the similarity of RNA-FM embeddings for miRNA sequences from the same family for miRNA families with more than 20 sequences. The five most abundant miRNA families were visualised in Supplementary Fig. S11a. We observed that miRNA sequences from the same family show significantly higher clustering based on average silhouette score when compared to randomly assigned sequence family (*p* < 0.0001) (Supplementary Fig. S11b). This analysis suggest that RNA-FM features can be beneficial in resolving substructures within RNA types.

We then evaluated whether deep learning embeddings could capture complex interpretable features. A linear regression model was trained to predict each continuous interpretable feature (Supplementary Fig. S12 and S13, Supplementary Table S4); whereas logistic regression models were trained to predict binary interpretable features (Supplementary Table S5). Compared to the highest absolute correlation coefficient squared (*r*^2^) between each interpretable feature and RNA-FM embedding dimensions, regression models were able to achieve R^2^ scores greater than 0.9 for most features; whereas only nucleotide U had highest absolute correlation coefficient squared exceeding 0.9 (*r*^2^ = 0.91) (Supplementary Fig. S12). For binary values, logistic regression models were able to achieve AUC scores of ∼0.99 (Supplementary Table S5). R^2^ score ranges of 0.51–0.99, compared to ranges of 0.04–0.91 from highest absolute correlation coefficient squared, confirms our previous observations that RNA-FM encode rich representations of RNA sequences across all embedding dimensions.

Similarly, we trained linear regression models to predict siRNA efficacy using Huesken (13) and Takayuki (14) datasets, separately, which showed that combining both interpretable features and selected RNA-FM features showed improved performance of ∼0.05 for Huesken dataset and ∼0.02 for Takayuki dataset compared to interpretable features only (Table 4). Due to the low sample size, we evaluated the prediction performance using 10-fold cross validation. This prediction performance supports RNA-FM’s potential in siRNA efficacy prediction even using a linear regression model.

**Table 4.** Prediction performance (R^2^ Score, mean ± standard deviation across 10 folds) of linear regression predicting siRNA efficacy from Huesken and Takayuki datasets, respectively, using 10-fold cross validation.

|  | Huesken Dataset | Takayuki Dataset |
| --- | --- | --- |
| Interpretable Features only | 0.233 ± 0.040 | 0.362 ± 0.128 |
| RNA-FM embeddings (selected dimensions only) and Interpretable Features combined | 0.283 ± 0.055 | 0.381 ± 0.096 |

### RNAExplorer, a web platform for small RNAs and their features

To support a wide range of scientists in analysing and comparing small RNA sequences, we developed RNAExplorer (www.rnaexplorer.com), a user-friendly web application that provides interactive access to RNA-FM representations and comparative feature analyses without requiring specialist computational expertise (Fig. 9). The interface integrates sequence input, feature selection and visualisation within a single view. RNAExplorer showcases two complementary embeddings: an RNA-FM embedding, which places user-provided sequences within the landscape of known and published small RNA classes, and an interpretable feature embedding, which represents sequences using biologically meaningful features such as length, GC content, mononucleotide composition, and k-mer frequencies. User-provided sequences are overlaid onto both embedding plots, enabling direct comparison with reference siRNA, miRNA, and piRNA sequences. Interactive controls, including sequence input, dataset colouring, and feature selection, allow users to dynamically explore class structure and sequence similarity. Individual sequences displayed in the plots are fully interactive. Selecting a point redirects the user to the original data source, providing access to related information.

**Figure 9.**
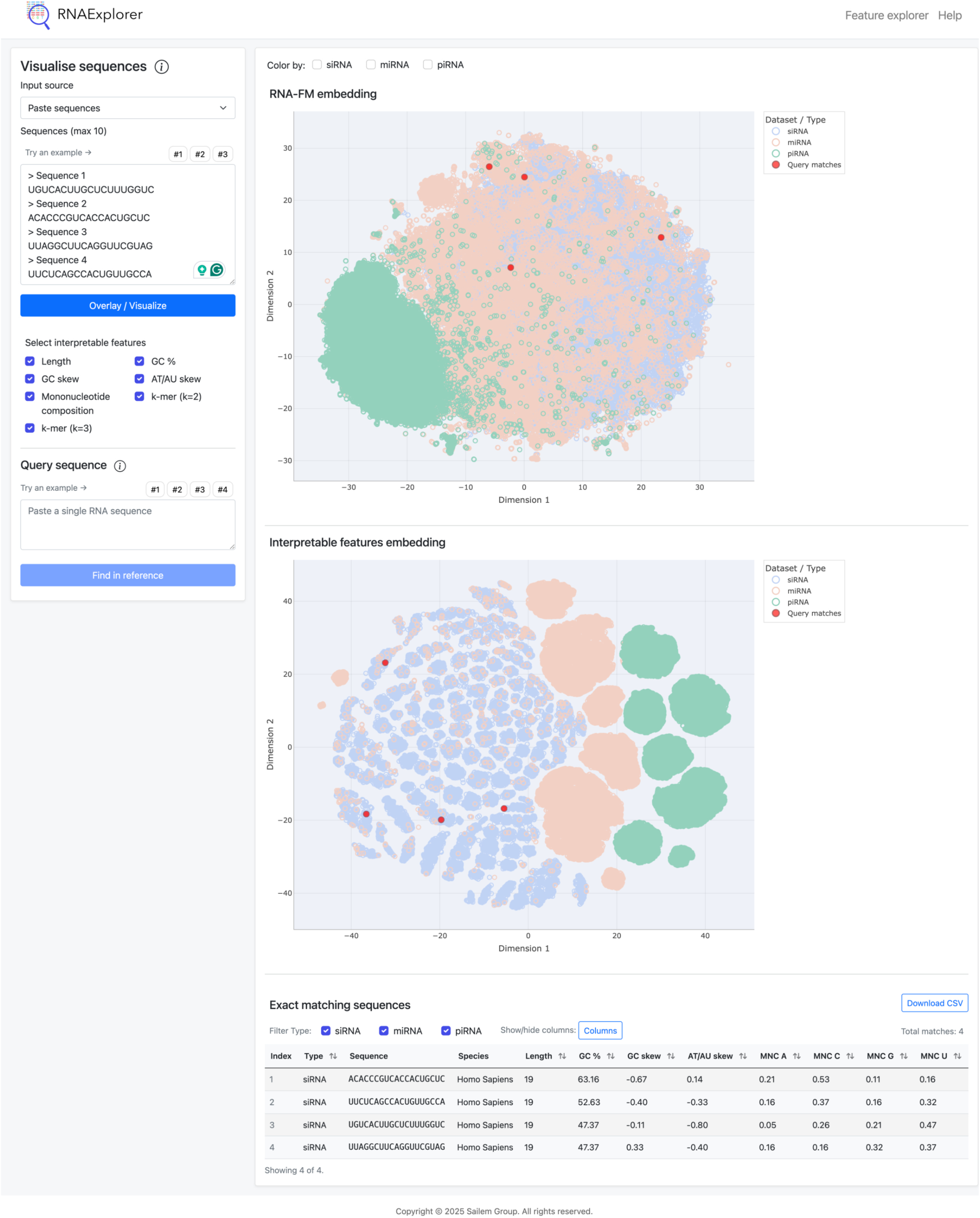
A screenshot of RNAExplorer web interface. Red dots in interpretable- and deep learning-based embeddings highlight four siRNA sequences designed to target the *SNCA* gene.

In addition to the embedding visualisations, RNAExplorer provides integrated tabular views to support detailed inspection and downstream analysis of sequences (Fig. 9). The interface retrieves most similar reference sequences based on deep learning features and exact matching sequences that contain the query sequence. Users can customise the table view by showing or hiding specific columns according to their analysis requirements and may download the displayed data for offline use. Species annotations for miRNA sequences are included for each reference sequence, with species names linked to their corresponding source database, facilitating biological interpretation.

While existing RNA analysis platforms provide individual sequence analysis and search functionalities, we did not identify a platform providing the representation-based capabilities implemented in RNAExplorer. Specifically, RNAExplorer integrates RNA-FM embedding generation and embedding-based similarity search with analysis and visualisation of deep-learning and interpretable feature spaces, enabling query sequences to be positioned and compared within a reference landscape of small RNAs (Table 5). These capabilities are further integrated with fragment-based sequence search and conventional sequence-property analyses, including GC content, nucleotide composition, and nucleotide skew calculations.

**Table 5.** Comparison of publicly available web applications for RNA sequence analysis against RNAExplorer across key functionalities.

| Functionalities | RNAExplorer<br>(Our web-based application) | RNAcentral<br>(24) | Rfam<br>(25,26) | EMBOSS<br>(27) | OligoPool<br>(28–30) | OligoCalc<br>(31) |
| --- | --- | --- | --- | --- | --- | --- |
| RNA-FM embedding generation | ✓ | × | × | × | × | × |
| RNA-FM embedding-based similarity search | ✓ | × | × | × | × | × |
| Visualisation in deep-learning feature space | ✓ | × | × | × | × | × |
| Visualisation in interpretable feature space | ✓ | × | × | × | × | × |
| Query placement within a reference small-RNA landscape | ✓ | × | × | × | × | × |
| Fragment-based sequence search | ✓ | ✓ | ✓, via RNAcentral | × | × | × |
| GC content calculation | ✓ | × | × | ✓, via geecee | ✓ | ✓ |
| Nucleotide composition calculation | ✓ | × | × | ✓, via compseq | × | × |
| Nucleotide skew calculation | ✓ | × | × | Not directly; can be derived using EMBOSS tools | × | × |

### RNAExplorer provides insights relevant to RNA design

From the embedding-based similarity search, we asked how similarity with existing sequences may provide insights to RNA sequence design, such as biological compatibility. To assess that, we measured the similarities between a given siRNA sequence with top *k* most similar miRNA sequences (*k* = 20, 50, or 100) and correlated the average similarity to its efficacy for sequences in both Huesken (13) and Takayuki (14) datasets (Supplementary Fig. S14). Across both datasets and all values of *k*, miRNA similarity showed a consistent but modest positive correlation with efficacy (r = 0.12–0.19; all *p* < 0.0001). Thus, siRNAs with greater similarity to endogenous miRNAs tended to have slightly higher efficacy. This suggests that higher similarity of siRNA candidates to miRNA sequences is associated with increased efficacy, which may potentially reflect higher biological compatibility.

Next, we illustrate the utility of RNAExplorer for guiding siRNA sequence design for *SNCA*, a gene whose dysregulation contributes to neuronal dysfunction and degeneration in Parkinson’s disease (32,33). A researcher evaluates four candidate siRNA sequences (Sequence 1: UGUCACUUGCUCUUUGGUC, Sequence 2: ACACCCGUCACCACUGCUC, Sequence 3: UUAGGCUUCAGGUUCGUAG, Sequence 4: UUCUCAGCCACUGUUGCCA) designed to target *SNCA* (Fig. 9). Computing the similarity of reference miRNAs to the candidates allows retrieval of the top *k* most similar miRNAs (*k* = 20, 50, or 100) for each candidate based on their distance to the query’s deep RNA-FM embeddings (Methods). Across four candidates, Sequences 1 and 4 have higher similarity across top 20 most similar miRNAs followed by Sequences 3 and 2 with decreasing similarity (Fig. 10a). Specifically Sequence 4 shows similarity to 6 endogenous miRNAs across different species including the human hsa-miR-7112-3p, followed by Sequence 1 with similarity to 3 miRNAs, then Sequences 2 and 3 (Fig. 10b and Supplementary Table S6). Sequence 4 may be especially relevant because of its similarity to hsa-miR-7112-3p, which has been reported to suppress PERK expression (34). PERK signalling is implicated in the endoplasmic reticulum stress response associated with α-synuclein accumulation resulting from *SNCA* dysfunction (35). Given the modest association observed between miRNA similarity and siRNA efficacy, these results prioritise Sequence 4, followed by Sequence 1, as candidates for further experimental evaluation, while providing an opportunity to investigate whether similarity to endogenous miRNAs is also related to biological compatibility. Together, these findings illustrate how RNAExplorer can identify potentially biologically relevant relationships between candidate siRNAs and endogenous RNAs that may inform candidate prioritisation, while further experimental validation is required to establish their functional significance.

**Figure 10.**
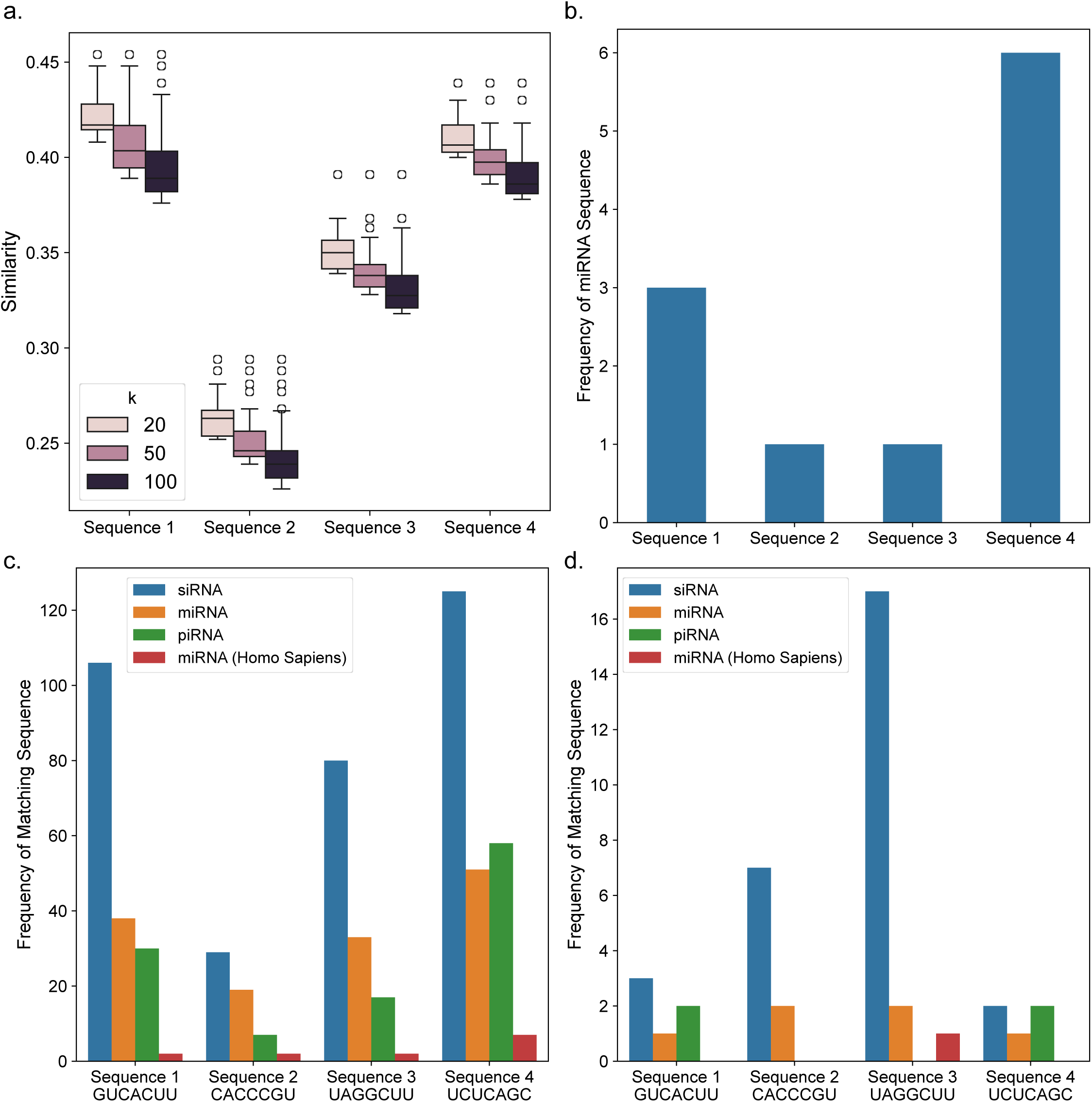
Similarity and seed analysis for four siRNA sequences targeting *SNCA* gene. (a) Similarity distribution for each sequence with top *k* most similar miRNA sequences, where *k* = 20, 50 and 100. (b) Number of miRNAs among top 20 most similar sequences. (c) Number of sequences containing the seed fragment across RNA classes, including siRNA (blue), miRNA (orange), piRNA (green), and Homo sapiens miRNA (red) shown as a separate category. (d) Number of sequences exhibiting an exact seed match at positions 2–8 for the same set of seed sequences and RNA types.

Beyond global sequence similarity, RNAExplorer enables systematic assessment of seed-mediated off-target potential. A major challenge in siRNA design lies in balancing on-target efficacy with off-target effects driven by miRNA-like seed interactions (nucleotides 2–8), which can result in repression of multiple transcripts (36). RNAExplorer allows users to search for a specified sequence fragment and identify all database sequences that contain it, retrieving the number of matches along with their RNA class and species of origin. The seed of the four sequences occurs in sequences with variable deep and interpretable features (Supplementary Fig. S15). The seed fragment of Sequence 3 matches the seed region at positions 2–8 of one human miRNA, which may indicate a greater potential for shared miRNA-like targeting and therefore merits further off-target assessment (Fig. 10c, d and Supplementary Table S7). In contrast, although the seed fragments of Sequences 1, 2 and 4 occur elsewhere within several human miRNA sequences, none aligns with the corresponding miRNA seed region at positions 2–8. These findings may suggest that Sequences 1, 2, and 4 may be more suitable candidates for mitigating potential off-target effects. Such comparisons allow the researcher to weigh trade-offs between biological similarity and potential off-target risk. Together, this example illustrates how RNAExplorer supports informed siRNA design by integrating deep learning-based similarity, interpretable sequence features, and potential off-target effects based on seed-level analysis.

## DISCUSSION

Therapeutic siRNA design is challenged by the need to balance efficacy, specificity and biological compatibility (37). Comparing synthetic siRNAs with endogenous small-RNA classes may provide a broader biological context for understanding sequence properties relevant to these challenges. Accordingly, the principal contribution of this study is a systematic, large-scale comparison of compositional, positional, structural and thermodynamic properties across siRNAs, miRNAs and piRNAs. We establish a quantitative reference landscape of these properties, determine how information associated with them is represented within RNA-FM embeddings, evaluate the predictive utility of these representations, and translate the resulting framework into RNAExplorer. By positioning siRNAs within the broader landscape of endogenous small RNAs, this framework supports biological interpretation, hypothesis generation and informed candidate prioritisation.

The comparative analysis revealed distinct quantitative feature landscapes across siRNAs, miRNAs and piRNAs. Although several individual relationships align with established biochemical principles and previously reported characteristics of specific RNA classes, their systematic evaluation within a common framework demonstrates how the magnitude and combination of these properties vary across the three classes. These differences were further supported by unsupervised analyses and by the ability to predict small-RNA class from RNA-FM embeddings generated from sequence alone, providing evidence that class-associated information is represented within the embedding space. The separation between piRNA and other RNA types based on higher thermodynamic stability can be partially explained by their longer sequences and composition characterised by higher GC content and melting temperature (Fig. 2 and Table 3). These factors are known to increase RNA duplex stability through base stacking and base pairing in nearest-neighbour models (38). The lower variation in Gibbs free energy and minimum free energy of siRNA could reflect design-selected properties. The intermediate position of miRNAs for several features aligns with their conserved hairpin processing and seed-based targeting, maintaining a characteristic structure without extreme composition shifts (39,40). In contrast, piRNAs show broader, long-tailed distributions across key features most clearly in ΔG and MFE, indicating greater sequence heterogeneity. Current studies note that piRNA pathways can operate with relaxed targeting rules during recognition of transposon transcripts, which is consistent with greater sequence diversity in this class (41,42).

The interpretability of RNA-FM features was thoroughly explored through the t-SNE projection and correlation analysis, verifying that biological attributes ranging from structural to thermodynamic parameters are encoded within deep learning embeddings. The t-SNE plot of RNA-FM embeddings demonstrated a clear separation of piRNAs from miRNAs and siRNAs, with the latter two forming overlapping regions. As t-SNE preserves local neighbourhoods but not global distances, conclusions about class differences were therefore evaluated using Principal Component Analysis (PCA). This confirmed that the observed separation was generally consistent across dimensionality-reduction methods. The tight clustering of piRNAs supports the idea that RNA-FM effectively captures their distinctive sequence characteristics, such as strong 5′-uridine bias, greater structural diversity, and longer sequence lengths. The transition region observed between siRNAs and miRNAs may reflect structural similarity, particularly for endogenous siRNAs that resemble miRNA precursors. Moreover, mapping interpretable features onto the RNA-FM embeddings revealed consistent spatial gradients in thermodynamic values, base composition, strand asymmetry, and position-specific nucleotide preferences. These embeddings, obtained without task-specific labels, demonstrate how foundation models can uncover latent biological structure in RNA data, offering a scalable and generalisable framework for interpreting RNA function and diversity.

Distinct correlation patterns along with biological feature predictions revealed that the deep learning model distinguishes between sequence-wide and position-specific information, as well as mononucleotide and dinucleotide information. Hierarchical clustering grouped interpretable properties (iCls) and RNA-FM dimensions (dCls) according to their correlation profiles, revealing how the two feature spaces co-vary and identifying relationships that are shared across, or specific to, different RNA classes. This mapping improves the biological interpretability of the embedding space and helps identify RNA-FM dimensions relevant to specific sequence properties for comparative and predictive analyses. However, because detailed functional annotations are unavailable for many of the analysed sequences, the biological meaning of these clusters remains difficult to determine.

Classification and feature prediction analyses increase confidence in the predictive utility of RNA representations. Similarly, higher clustering quality across RNA types and across abundant miRNA families than random baselines also suggest RNA-FM captures type-specific information and finer-grained subgroups within RNA types. Such insight not only increases transparency but also demonstrates their utility for downstream applications, enabling users to select or constrain features based on biological relevance rather than treating the model as a black box.

RNAExplorer web application enables researchers to interrogate their sequences in the context of other small RNAs, providing insights into similarity, potential off-target effects, and possible functional roles. As such, it empowers researchers to obtain rapid, interpretable insights into their own sequences and map them to sequences in our database. This can be particularly useful for designing and prioritising siRNA sequences by gaining insights into similar miRNA and piRNA sequences or by identifying fewer human miRNA seed matches. This can also facilitate generating hypotheses on potential functions for a given small RNA by examining in greater detail the functional roles of matched or similar miRNAs. RNAExplorer therefore supports comparative analysis and candidate prioritisation, while experimental validation remains necessary to establish gene-silencing efficacy and specificity.

A major challenge in studying small RNA sequences is that our knowledge of miRNA and piRNA biological functions is incomplete, which limits the functional interrogation of these features. For siRNAs, assessing biological compatibility is particularly challenging because in vivo validation is constrained by ethical and practical considerations, while sufficiently large and well-annotated datasets of experimentally measured siRNA efficacy remain limited. In our analysis, similarity to endogenous miRNAs showed a modest association with siRNA efficacy, providing preliminary evidence that similarity captured by RNA-FM embeddings may reflect information relevant to siRNA activity. However, efficacy represents only one aspect of biological compatibility and cannot fully capture properties such as off-target effects, toxicity, or in vivo tolerability. The modest observed correlation may also reflect the limited size and scope of the available efficacy dataset, and therefore requires further investigation using larger, well-annotated datasets and, where feasible, experimental validation of functional and biocompatibility-related outcomes. Finally, this study focused exclusively on three well-characterised small RNA types. Going forward, it would be interesting to integrate other RNA classes such as tRNA-derived fragments, snoRNAs, or lncRNA-derived small RNAs.

In future, our comparative framework could be generalised to include diverse taxonomic groups or developmental contexts, enabling a systems-level view of small RNA evolution and specialisation across species or tissues. By integrating deep learning features, scientists can select RNA-FM features that are strongly correlated to interpretable features and gain quick insights into their data. RNA-FM’s interpretability helps to understand the success of recent approaches that utilised foundation models to perform downstream tasks such as siRNA efficacy prediction (20) and ncRNA-protein interaction classifier (23). These insights can be extended to improve the classification of newly discovered or poorly annotated small RNAs and may help in identifying functionally relevant subclasses with distinct regulatory roles in development, immunity, or disease.

This systematic comparison of small RNAs provides a framework for investigating relationships between sequence properties, RNA-FM representations and RNA function, with potential relevance to therapeutic RNA research. RNAExplorer enables researchers to obtain interpretable features and RNA-FM representations of their sequences for comparative analyses and downstream applications, including candidate prioritisation. By consolidating the distinct characteristics of major small-RNA classes, our study provides a foundation for exploring how these properties relate to function and may inform future therapeutic RNA design.

## DATA AVAILABILITY

The raw sequence datasets used in this study are publicly available from established repositories. The miRNA sequences were retrieved from miRBase https://www.mirbase.org/download/, and piRNA sequences were accessed via DASHR2 https://dashr2.lisanwanglab.org/. Positional features such as U at position 1, G at position 1, UU across sequence, etc., were computed using the code available at https://doi.org/10.1093/bioinformatics/btae577. Sequence-level metadata and feature matrices are provided in tab-delimited format. All extracted deep and interpretable features are available from RNAExplorer repository (https://github.com/sailem-group/RNAExplorer).

## CODE AVAILABILITY

The source code for RNAExplorer and a Jupyter notebook for reproducibility are available on https://github.com/sailem-group/RNAExplorer. The repository includes the setup instructions and the code required to deploy and run the RNAExplorer web application. The Jupyter notebook contains all steps required to reproduce the analysis results and figures, ranging from processing to visualisation.

## Acknowledgements

We acknowledge all members of the TransNat consortium, particularly its lead, Matthew Wood, and the Sailem Group for useful discussions.

## Contributions

H.S. conceived and designed the study. S.J. and C.O. conducted the literature review, curated the sequence data, and performed data analysis. H.S. and S.J. designed and developed the RNAExplorer application. H.S., S.J., and C.O. wrote the paper. All authors have read and approved the manuscript.

## Funding

This work is funded by the MRC TransNat grant (MR/X008029/1).

## Notes

### Competing Interest Statement

The authors have declared no competing interest.

### Summary of Updates

All sections and figures are updated to reflect revised data analysis. The discussion is expanded.

